# Loss of Depalmitoylation Exaggerates Synaptic Upscaling and Leads to Neuroinflammation in a Lysosomal Storage Disease

**DOI:** 10.1101/2021.12.16.473002

**Authors:** Kevin P Koster, Eden Flores-Barrera, Emilce Artur de la Villarmois, Thu T. A. Nguyen, Amanda Niqula, Lorena Y Noriega-González, Zach Fyke, Adriana Caballero, Stephanie M. Cologna, Kuei Y. Tseng, Akira Yoshii

## Abstract

Palmitoylation and depalmitoylation are the dichotomic processes of lipid modification regulating protein trafficking, recycling, and degradation, thereby controlling proteostasis. Despite our understanding of palmitoylation, depalmitoylation is far less studied. Here, we study a lysosomal depalmitoylating enzyme, palmitoyl-protein thioesterase 1 (PPT1), associated with the devastating neurodegenerative condition CLN1 disease and show that dark-rearing *Ppt1^-/-^* mice, which induces synaptic upscaling *in vivo*, worsen the symptoms. In *Ppt1^-/-^* cortical neurons, upscaling induction triggers exaggerated responses of synaptic calcium-permeable AMPA receptors composed of palmitoylated GluA1 subunits. Consequently, *Ppt1^-/-^* visual cortex exhibits hypersynchrony *in vivo*. Remarkably, we also find an overload of palmitoylated A-kinase anchor protein 5 (Akap5) in *Ppt1^-/-^* mouse brains, leading to microglial activation through NFAT. These findings indicate Ppt1 acts as a gatekeeper of homeostatic plasticity by regulating the proteostasis of palmitoylated synaptic proteins. Moreover, our results suggest that perturbed depalmitoylation results in neuroinflammation, which is common to neurodegenerative diseases.

## Introduction

Protein homeostasis, or proteostasis, is a fundamental molecular mechanism whereby quality, quantity, and distribution of proteins are rigorously controlled through synthesis, folding, trafficking, and degradation. Proteostasis is germane to neurodegeneration and lysosomal storage diseases, as well as cancer, diabetes, and normal aging (Agyemang et al., 2015; Drie, 2011; Hetz and Glimcher, 2011; Labbadia and Morimoto, 2014). This dynamic regulation of proteins depends on the balance between supply and breakdown. Several mechanisms carry out protein degradation, including the unfolded protein response (Cao and Kaufman, 2012; Walter and Ron, 2011), ubiquitination-mediated proteasomal degradation (Nandi et al., 2006), autophagy (Nixon, 2013; Saha et al., 2018), and lysosomal degradation (Lie and Nixon, 2018; Xu and Ren, 2015). The lysosome in particular plays a critical role in digesting proteins modified with glycosyl or lipid chains. Accordingly, there are over 50 lysosomal storage disorders, and most of them are associated with mutations in lysosomal enzymes that detach protein modifications (Platt et al., 2018).

Infantile neuronal ceroid lipofuscinosis (CLN1) is a devastating pediatric neurodegenerative disease and is classified as a lysosomal storage disease (Nita et al., 2016). Children afflicted by CLN1 show no detectable symptoms during the first 6 to 12 months of life. Following this seemingly typical developmental period, CLN1 patients present with a deceleration of head growth (microcephaly), loss of vision, motor deterioration, and epilepsy that constitute a severe developmental regression and result in premature death by age 5 (Haltia, 2003; Nita et al., 2016). CLN1 is caused by mutations in the gene *CLN1*, which encodes the enzyme palmitoyl-protein- thioesterase (Ppt1) (Vesa et al., 1995).

Ppt1 is a depalmitoylating enzyme, which removes the 16-carbon fatty acid, palmitate, from postranslationally palmitoylated proteins, thereby regulating their trafficking, localization, and turnover. Appreciation for the functional importance of protein palmitoylation at the synapse has increased substantially in recent years (Fukata and Fukata, 2010). On the other hand, the significance of depalmitoylation in synaptic plasticity or neuronal development remains less clear. Early studies of Ppt1 function demonstrated its activity toward signaling molecules, such as H-Ras (Camp and Hofmann, 1993), and demonstrated in non-neuronal cell types that it is localized to the lysosome (Verkruyse and Hofmann, 1996). Accordingly, loss of PPT1 in CLN1 disease causes the robust lysosomal accumulation of autofluorescent lysosomal storage material (AFSM) frequently referred to as lipofuscin. However, Vesa and colleagues first suggested that Ppt1 is also secreted (Vesa et al., 1995), and additional evidence showed that Ppt1 localizes beyond the lysosome to the broader endolysosomal compartment, including synaptic vesicles (Ahtiainen et al., 2003, 2006; Lehtovirta et al., 2001; Lyly et al., 2007). Further, a recent study probing for Ppt1 substrates using a stringent proteomic screen corroborated that a portion of PPT1 is secreted to the synaptic cleft and showed that roughly 10% of the synaptic palmitoyl proteome (palmitome), representing over 100 proteins, is depalmitoylated by Ppt1. These findings clarified that synaptic proteins, especially transmembrane proteins, make up a large proportion of Ppt1 substrates. Interestingly, this study identified the α-amino-3-hydroxy-5-methyl-4-isoxazolepropionic acid receptor (AMPAR) subunit GluA1 as a substrate (Gorenberg et al., 2020).

Given the growing evidence for Ppt1 as a synaptic enzyme, defining its role in synaptic plasticity is essential to deciphering CLN1 disease pathophysiology and has broader implications for cortical circuit development. Two major forms of synaptic plasticity, Hebbian and homeostatic, govern brain circuit formation and function (Espinosa and Stryker, 2012; Keck et al., 2017; Lau and Zukin, 2007; Pozo and Goda, 2010; Turrigiano, 2017). Hebbian plasticity is a conceptual framework where synapses that “fire together, wire together” (Katz and Shatz, 1996), such that local synaptic strength is either strengthened or weakened depending on the frequency or intensity of the incoming stimulus. High frequency stimuli induce synaptic strengthening in a process called long-term potentiation (LTP), while low frequency stimuli triggers synapse weakening termed long-term depression (LTD). One study shows that loss of Ppt1 suppresses LTP in *ex vivo* hippocampal slice recordings (Sapir et al., 2019).

Synaptic scaling, on the other hand, is a form of homeostatic plasticity by which neurons compensate for chronic changes in network activity by increasing or decreasing the number of receptors on the synaptic surface (O’Brien et al., 1998; Turrigiano et al., 1998). Explicitly, in response to reduced chronic activity, AMPARs are inserted into the membrane (synaptic upscaling) while, in response to a chronic increase in afferent activity, AMPARs are endocytosed (synaptic downscaling) (O’Brien et al., 1998; Turrigiano et al., 1998). Detailed analyses have revealed that synaptic upscaling is largely constituted by increases in calcium-permeable AMPARs (CP-AMPARs)(Ju et al., 2004; Lee, 2012; Thiagarajan et al., 2005), typically composed of GluA1 homomers (Burnashev et al., 1992; Wenthold et al., 1996). Importantly, synaptic scaling is disrupted in neurodevelopmental and neurodegenerative diseases, such as Rett syndrome and Alzheimer disease (Blackman et al., 2012; Gilbert et al., 2017; Pratt et al., 2011; Qiu et al., 2012; Styr and Slutsky, 2018). Moreover, CP-AMPARs are implicated in epilepsy, stroke, and motor neuron disease (Netzahualcoyotzi and Tapia, 2015; Noh et al., 2005; Quintana et al., 2015; Rajasekaran et al., 2012; Yamashita and Kwak, 2018). However, the link between lysosomal depalmitoylation and homeostatic plasticity mechanisms has not been studied. Therefore, we sought to determine if synaptic scaling is dysregulated in *Ppt1^-/-^* mice and if the abnormal response contributes to CLN1 pathology.

Here, we describe the mechanism by which synaptic scaling is dysregulated in *Ppt1^-/-^* neurons. We find that synaptic upscaling of CP-AMPARs is exaggerated in *Ppt1^-/-^* neurons that correlates with an overload of palmitoylated GluA1. Further, we show that dark rearing (dark reared/rearing; DR) *Ppt1^-/-^* mice, an *in vivo* model of synaptic upscaling in the visual cortex, accelerates neuroinflammation and brain atrophy. Notably, A-kinase anchor protein 5 (Akap5) and its associated signaling pathways contribute to CLN1 pathology through the hyperactivation of nuclear factor activated in T-cells (NFATc3) and neuroinflammation, representing a link between aberrant synaptic plasticity and an inflammatory cascade in CLN1 and opening a new set of therapeutic targets. Together, these data indicate that Ppt1 regulates proteostasis of CP-AMPARs through depalmitoylation and plays a critical role in controlling homeostatic plasticity. Further, our findings shed light on how loss of Ppt1 function causes hypersynchrony and neurodegeneration of neural circuitry.

## Results

The *Ppt1^-/-^* mouse model develops detectable myoclonic jerks and motor dysfunction by four months of age (Gupta et al., 2001). However, considering the early timeline of disease in humans and developmental regulation of Ppt1 in rodents (Koster et al., 2019; Suopanki et al., 1999a, 1999b, 2000), we anticipate, and have previously demonstrated (Koster et al., 2019), that disruption of developmental synaptic plasticity mechanisms is germane to disease pathogenesis. Therefore, in the current study we examine AMPAR plasticity at timepoints spanning the first 10 weeks postnatal in presymptomatic *Ppt1^-/-^* mice.

There are two major advantages of focusing on the mouse visual system to study the role of PPT1 in synaptic plasticity. First, patients with CLN1 suffer progressive blindness that involves retinal, thalamic, and cortical degeneration, which is preceded by epileptic activity (Augustine et al., 2021; Koster and Yoshii, 2019; Nita et al., 2016). Thus, the visual cortex of *Ppt1^-/-^* mice is a disease-relevant brain region. Second, critical period plasticity has been extensively studied in rodent visual cortex and follows two well-characterized rules, Hebbian and homeostatic plasticity (Keck et al., 2017; Turrigiano and Nelson, 2004). Therefore, there are several established experimental paradigms to study experience-dependent plasticity in the central visual system including monocular deprivation and DR.

### Dark rearing *Ppt1^-/-^* mice Exacerbates Disease Pathology in the Visual Cortex

Previous studies demonstrated that DR mitigated disease progression in mouse models of other neurodevelopmental disorders with some overlapping symptoms to CLN1, including Rett syndrome (Durand et al., 2012) and Angelman syndrome (Yashiro et al., 2009). Therefore, we followed this approach, seeking to determine if DR would alleviate disease progression in *Ppt1^-/-^* mice. We reared groups of wild-type (WT) and *Ppt1^-/-^* mice in complete darkness beginning at postnatal day (P) 10-12, before pups open their eyes, and sacrificed animals for analysis of common disease features of CLN1, such as autofluorescent storage material (AFSM) and decreased cortical thickness, as well as mortality.

Contrary to the finding that DR alleviated symptoms in *Mecp2^-/-^* or *Ube3a^m-/p+^* mice (Durand et al., 2012; Yashiro et al., 2009), we found that it instead exacerbated disease symptoms in the visual cortex of *Ppt1^-/-^* mice (**Figure 1**). The hallmark of CLN1, AFSM, increased in the visual cortex (**Figure 1A-B**) and cortical thickness decreased in DR-*Ppt1^-/-^* mice compared to WT or *Ppt1^-/-^* animals raised under 12-hr dark and light cycle (light reared; LR) (**Figure 1C**), demonstrating an accelerated cortical atrophy in DR animals. Importantly, mortality occurred significantly earlier as a result of severe seizures in DR-*Ppt1^-/-^* animals (**Figure 1D**). These results indicate that AFSM accumulation is activity-dependent in DR-*Ppt1^-/-^* mice and suggest that lack of lysosomal depalmitoylation impairs proteostasis of palmitoylated proteins, interfering with synaptic plasticity before brain atrophy occurs.

**Figure 1.**
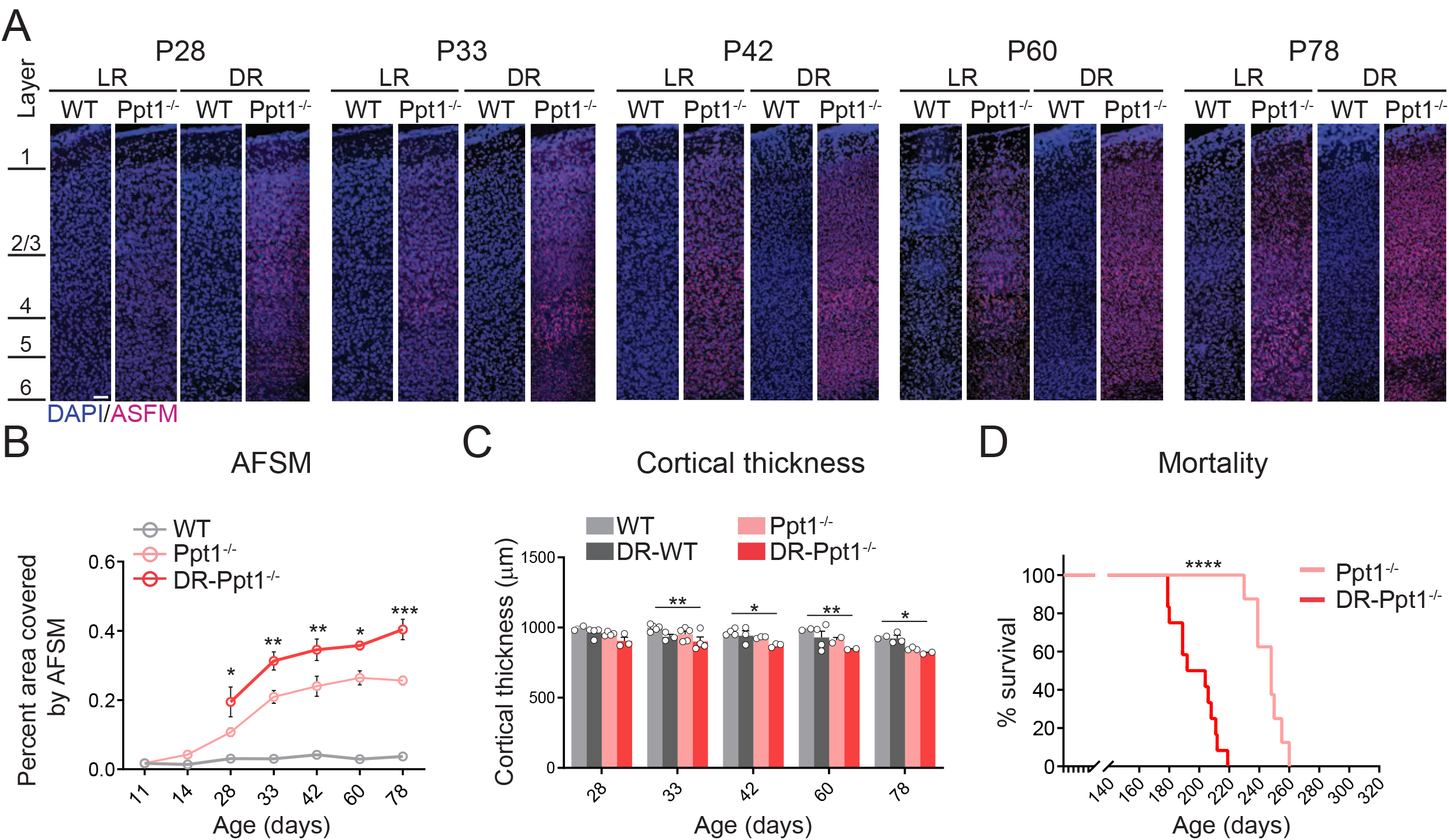
Dark rearing *Ppt1^-/-^* animals exacerbates disease pathology. **(A)** Representative sagittal sections of medial visual cortex from LR-WT, LR-*Ppt1^-/-^*, DR-WT, and DR-*Ppt1^-/-^* mice showing the accumulation of ASFM with age (postnatal ages 28-78). Scale bar = 50μm. **(B)** Quantification of the percent area covered by AFSM. Two-way ANOVA: interaction condition (genotype and rearing) x age (F (8, 41) = 3.072, ** P=0.0084); main effect of genotype/rearing (F (2, 41) = 174.5, **** P<0.0001); main effect of age (F (4, 41) = 13.23, ****P<0.0001). Tukey’s multiple comparison indicated on graph: *p=0.0157 P28 LR-*Ppt1^-/-^* vs. P28 DR-*Ppt1^-/-^*; **p=0.0034 P33 LR-*Ppt1^-/-^* vs. P33 DR- *Ppt1^-/-^*; **p= 0.0030 P42 LR-*Ppt1^-/-^* vs. P42 DR-*Ppt1^-/-^*; *p=0.0375 P60 LR-*Ppt1^-/-^* vs. P60 DR-*Ppt1^-/-^*. ***p= 0.0007 P78 LR-*Ppt1^-/-^* vs. P78 DR-*Ppt1^-/-^*. Data represent mean ± SEM. N=3-4 animals/group. **(C)** Quantification of cortical thickness across age in WT, DR-WT, *Ppt1^-/-^,* and DR-*Ppt1^-/-^* mice. Two-way ANOVA: no interaction of condition (genotype and rearing) x age (F (12, 43) = 0.4683, P=0.9223); main effect of genotype/rearing (F (3, 43) = 14.96, ****P<0.0001); main effect of age (F (4, 43) = 5.330, **P=0.0014). Tukey’s multiple comparison (simple effect within age) indicated on graph: **p=0.0046 P33 LR-WT vs. P33 DR-*Ppt1^-/-^*; *p= 0.0241 P42 LR-WT vs. P42 DR-*Ppt1^-/-^*; **p= 0.0078 P60 LR-WT vs. P60 DR-*Ppt1^-/-^*; *p= 0.0466 P78 LR-WT vs. P78 DR-*Ppt1^-/-^*; *p= 0.0401 P78 DR-WT vs. P78 DR-*Ppt1^-/-^*. Number of animals for each group (N=2-5) is displayed on the graph (individual points). Data represent mean ± SEM. **(D)** Kaplan-Meier plot of mortality in LR-*Ppt1^-/-^* and DR-*Ppt1^-/-^* mice. Log-rank (Mantel-Cox) test: ****p<0.0001 LR-*Ppt1^-/-^* vs. DR-*Ppt1^-/-^*. N=6 *Ppt1^-/-^*, 10 for DR-*Ppt1^-/-^*.

Disrupted lysosomal depalmitoylation or degradation is expected to result in overrepresentation of synaptic proteins on the cell surface through dysproteostasis. Considering DR only worsened the disease symptoms in *Ppt1^-/-^* animals and, further, that DR induces synaptic upscaling of AMPARs in the WT visual cortex (Desai et al., 2002a; Goel et al., 2006a; Ju et al., 2004; Shepherd et al., 2006; Thiagarajan et al., 2005), we reasoned that loss of Ppt1-mediated depalmitoylation causes a dysregulation of synaptic scaling (or related underlying molecular mechanism) that was exacerbated by DR. On the other hand, alleviation of disease symptoms by DR in the Rett syndrome mouse model was largely attributable to a correction of cortical inhibitory connectivity (Durand et al., 2012), implying GABAergic transmission may instead be affected in DR-*Ppt1^-/-^* mice. An analysis of cortical excitatory and inhibitory activity has not been studied in presymptomatic *Ppt1^-/-^* animals. Therefore, we next determined if glutamatergic or GABAergic circuitry was disrupted in the developing *Ppt1^-/-^* brain.

### Excitatory Synapse Formation Is Impaired in *Ppt1^-/-^* Visual Cortex during Early Development

To examine the role of Ppt1 in the developmental regulation of cortical synaptic activity, we conducted *ex-vivo* electrophysiological recordings from layer 2/3 visual cortical pyramidal neurons using an aCSF free of glutamate and GABA blockers, and a low-chloride-based internal solution to enable concurrent acquisition of excitatory (AMPAR) and inhibitory (GABAA) synaptic currents at a single-cell level as previously described (Flores-Barrera et al., 2017). At P14-15, both WT and *Ppt1^-/-^* mice showed similar levels of excitatory and inhibitory transmission onto layer 2/3 pyramidal neurons (**Figure 2A**). However, the frequency of AMPAR-mediated synaptic currents was markedly and preferentially reduced in the visual cortex of *Ppt1^-/-^* mice at P28-30 (**Figure 2B-C**), a disruption that was not associated with changes in the mean amplitude of the excitatory events (11.8 ± 0.9 pA in WT vs. 10.7 ± 0.5 pA in *Ppt1^-/-^*, p=0.3, unpaired student t-test). These data diverge from the finding in *Mecp2*^-/-^ mice (Rett syndrome), which exhibit inhibitory hyperconnectivity (Durand et al., 2012). Collectively, it is conceivable that the maturation of AMPAR-mediated excitatory synapses in the visual cortex is selectively impaired in the *Ppt1^-/-^* visual cortex.

**Figure 2.**
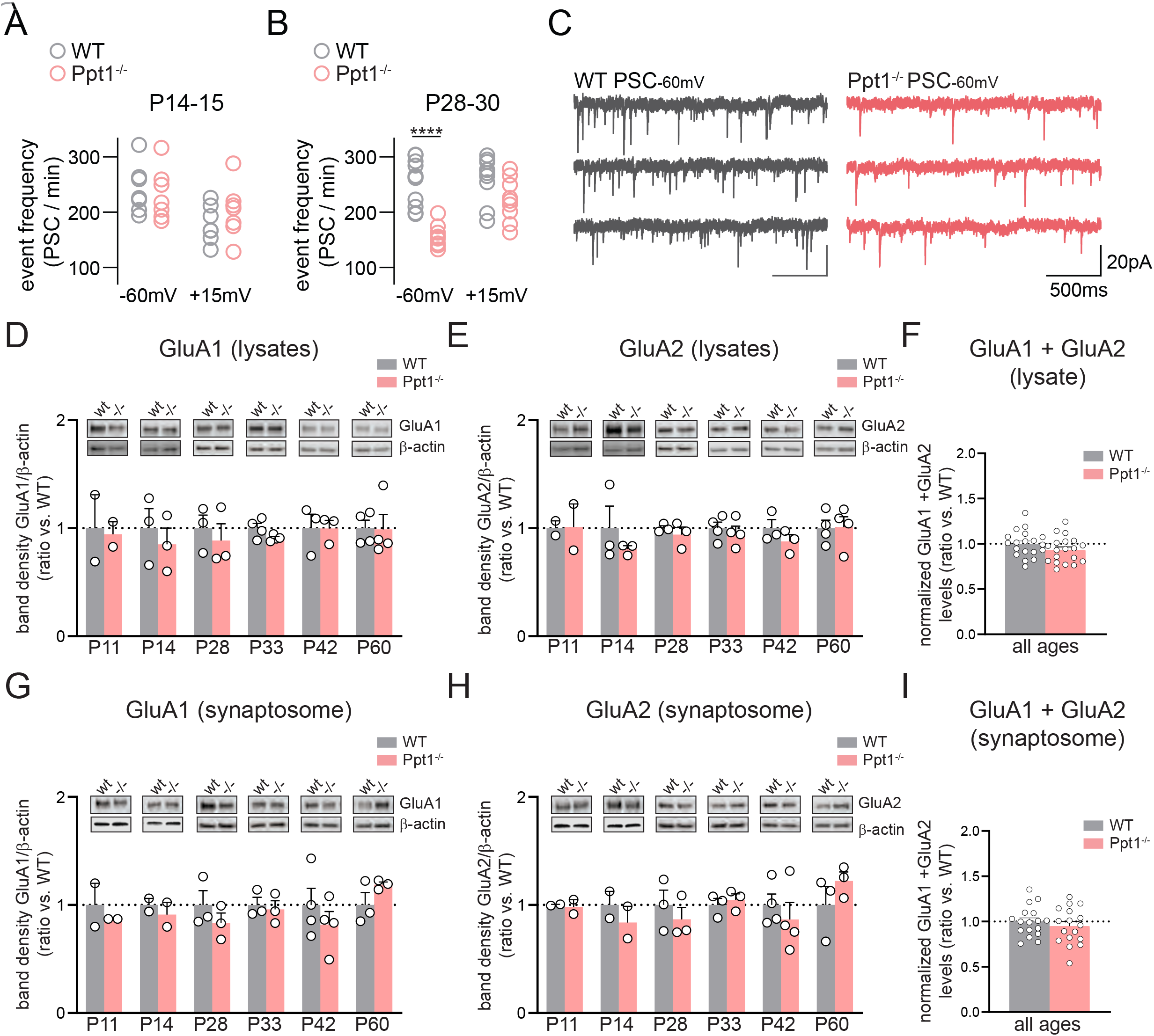
Developmental AMPA receptor transmission is decreased in *Ppt1^-/-^* visual cortex. **(A)** Quantification of sEPSC (−60mV) and sIPSC (+15mV) frequency from layer 2/3 visual cortical neurons in WT and *Ppt1^-/-^* mice at P14-15. n=6-7 cells, 3-4 animals/group. **(B)** Quantification of sEPSC (−60mV) and sIPSC (+15mV) frequency from layer 2/3 visual cortical neurons in WT and *Ppt1^-/-^* mice at P28-30. t-test: ****p<0.0001. n=9-11 cells, 3-4 animals/group, Data represent mean ± SEM. **(C)** Representative voltage-clamp recordings of sIPSCs and sEPSCs from layer 2/3 visual cortical neurons. **(D)** Representative immunoblots and quantification of total GluA1 levels in lysates from WT and *Ppt1^-/-^* visual cortices for ages P11-60. *Ppt1^-/-^* data are normalized to the WT mean value at each age and comparisons were made within age. N=2-4, Data represent mean ± SEM. **(E)** Representative immunoblots and quantification of total GluA2 levels in lysates from WT and *Ppt1^-/-^* visual cortices for ages P11-60. *Ppt1^-/-^* data are normalized to the WT mean value at each age and comparisons were made within age. N=2-4, Data represent mean ± SEM. **(F)** Quantification of the combined average of GluA1 and GluA2 levels in each occipital cortex lysate from WT and *Ppt1^-/-^* visual cortices for all ages examined. n= 19/group **(G)** Representative immunoblots and quantification of GluA1 levels in synaptosomes from WT and *Ppt1^-/-^* visual cortices for ages P11-60. *Ppt1^-/-^* data are normalized to the WT mean value at each age and comparisons were made within age. N=2-4, Data represent mean ± SEM. **(H)** Representative immunoblots and quantification of GluA2 levels in synaptosomes from WT and *Ppt1^-/-^* visual cortices for ages P11-60. *Ppt1^-/-^* data are normalized to the WT mean value at each age and comparisons were made within age. N=2-4, Data represent mean ± SEM. **(I)** Quantification of the combined average of GluA1 and GluA2 levels in each occipital cortex synaptosome from WT and *Ppt1^-/-^* visual cortices for all ages examined. n= 17/group

To determine if the reduced sEPSC frequency was due to a reduction in the AMPAR subunit content at *Ppt1^-/-^* synapses, we next performed immunoblotting for the GluA1 and GluA2 subunits in both lysates and synaptosomal fractions from WT and *Ppt1^-/-^* visual cortices. Total levels of GluA1 and GluA2 individually were unchanged throughout development (P11, 14, 28, 33, 42, 60), both in lysates (**Figure 2D-F**) and synaptosomes (**Figure 2G-I**). When combined within each fraction, levels of GluA1 and GluA2 at all ages did not demonstrate any change in the total AMPAR content in *Ppt1^-/-^* visual cortex (**Figure 2F, I**).

These results suggest that the activity-dependent turnover of AMPARs typical of excitatory synapse maturation was preferentially disrupted in *Ppt1^-/-^* mouse layer 2/3 pyramidal cells. We previously examined layer 2/3 pyramidal neuron dendrites in *Ppt1^-/-^* mice and found an increased number of dendritic spines with thin, immature morphology (Koster et al., 2019). Given the reduction of AMPAR sEPSC frequency in the visual cortex of *Ppt1^-/-^* mice but no change in the GluA levels or sEPSC amplitude, we reasoned that the fewer number of mature synapses are saturated with AMPARs, while extrasynaptic AMPARs are abundant due to deficient depalmitoylation. This notion led us to directly test the role of Ppt1 in synaptic upscaling of AMPARs. As several studies specifically implicate GluA1-containing AMPARs in synaptic scaling (Ju et al., 2004; Lee, 2012; Thiagarajan et al., 2005) and recent evidence points to a role for Ppt1 in depalmitoylating this AMPAR subunit (Gorenberg et al., 2020), we focused our analyses on the regulation of GluA1 in *Ppt1^-/-^* neurons.

### Synaptic Upscaling of GluA1 Is Exaggerated in *Ppt1^-/-^* Neurons *in vitro*

To determine whether Ppt1 regulates synaptic scaling, we examined this form of plasticity *in vitro* by incubating WT or *Ppt1^-/-^* primary cortical neurons in either bicuculline (20μm in DMSO) to induce downscaling, TTX (1μM in water) to induce upscaling, or vehicle control (VC, DMSO to a final concentration of 0.1%) for 48 hours, as previously reported (Diering et al., 2014; Sanderson et al., 2018; Shepherd et al., 2006; Turrigiano et al., 1998) (**Figure 3**).

**Figure 3.**
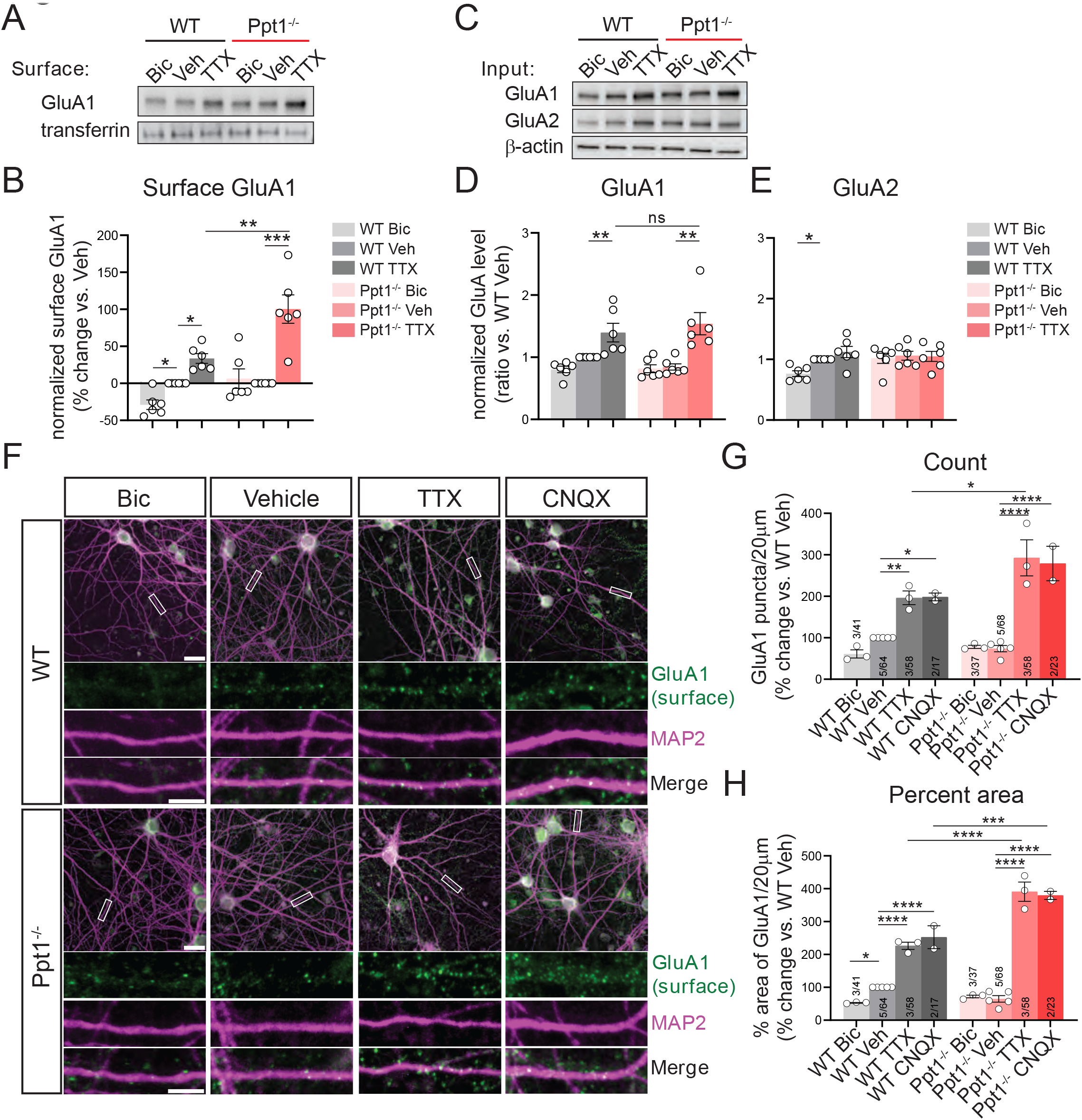
Synaptic scaling is exaggerated in primary cortical *Ppt1^-/-^* neurons. **(A)** Representative immunoblot of surface GluA1 and transferrin control following surface biotinylation assay in DIV16 cortical neuron cultures. Cells were treated with bicuculline (20μM), vehicle (DMSO, final concentration 0.1%), or TTX (1μM) for 48h. **(B)** Quantification of the surface GluA1 levels in WT and *Ppt1^-/-^* treated and untreated cortical neurons, normalized to transferrin for each experiment and expressed as percent change vs. vehicle condition. N=6 independent cultures. Two-way ANOVA: interaction genotype x treatment (F (2, 10) = 7.061, *P=0.0122); main effect of genotype (F (1, 5) = 14.21, *P=0.0130); main effect of treatment (F (2, 10) = 34.77, ****P<0.0001). Tukey’s multiple comparison indicated on graph: *p=0.0418 WT Bic vs. WT Veh; *p=0.0240 WT Veh vs. WT TTX; ***p=0.0002 *Ppt1^-/-^* Veh vs. *Ppt1^-/-^* TTX; **p=0.0038 *Ppt1^-/-^* TTX vs. WT TTX. Data represent mean ± SEM. **(C)** Representative immunoblots of GluA1, GluA2, and β-actin loading control for input samples (lysates). N=6 cultures. **(D)** Quantification of input GluA1 levels in WT and *Ppt1^-/-^* treated and untreated cortical neurons, normalized to β-actin for each experiment and expressed as percent change vs. vehicle condition. N=6 independent cultures. Two-way ANOVA: no interaction genotype x treatment (F (2, 10) = 2.902, P=0.1014); no main effect of genotype (F (1, 5) = 2.432e-005, P=0.9963); main effect of treatment (F (2, 10) = 14.81, **P=0.0010). Tukey’s multiple comparison indicated on graph: *p= 0.0106 WT Veh vs. WT TTX; ***p= 0.0001 *Ppt1^-/-^* Veh vs. *Ppt1^-/-^* TTX. Data represent mean ± SEM. **(E)** Quantification of input GluA2 levels in WT and *Ppt1^-/-^* treated and untreated cortical neurons, normalized to β-actin for each experiment and expressed as percent change vs. vehicle condition. N=6 independent cultures. Two-way ANOVA: interaction genotype x treatment (F (2, 10) = 9.617, **P=0.0047); no main effect of genotype (F (1, 5) = 1.699, P=0.2492); no main effect of treatment (F (2, 10) = 3.316, P=0.0786). Tukey’s multiple comparison indicated on graph: *p=0.0112 WT Bic vs. WT Veh. Data represent mean ± SEM. **(F)** Representative images of surface immunolabeled GluA1 in DIV16 WT and *Ppt1^-/-^* cortical neurons, treated with bicuculline (20μM), vehicle control (DMSO, final concentration 0.1%), or TTX (1μM) for 48h. Scale bars for low magnification images = 20μm, for high magnification = 5μm. **(G)** Quantification of the number of surface immunolabeled GluA1 puncta over 20μm of dendrite at a standard distance 50-100μm from the soma. N = 2-5 cultures/group (number of cultures and total number of cells analyzed are listed for each group on the graph). Data are normalized to the WT vehicle control (Veh) condition. Two-way ANOVA: interaction genotype x treatment (F (3, 18) = 5.620, **P=0.0067); main effect of genotype (F (1, 18) = 10.06, **P=0.0053); main effect of treatment (F (3, 18) = 52.74, ****P<0.0001). Tukey’s multiple comparison indicated on graph: **p= 0.0097 WT Veh vs. WT TTX; *p=0.0218 WT Veh vs. WT CNQX; ****p<0.0001 *Ppt1^-/-^* Veh vs. *Ppt1^-/-^* TTX; ****p<0.0001 *Ppt1^-/-^* Veh vs. *Ppt1^-/-^* CNQX; *p=0.0218 WT TTX vs. *Ppt1^-/-^* TTX. Data represent mean ± SEM. **(H)** Quantification of the percent area covered by surface immunolabeled GluA1 puncta over 20μm of dendrite at a standard distance of 50-100μm from the soma. N = 2-5 cultures/group (number of cultures and total number of cells analyzed are listed for each group on the graph). Data are normalized to the WT vehicle control condition. Two-way ANOVA: interaction genotype x treatment (F (3, 18) = 26.21, ****P<0.0001); main effect of genotype (F (1, 18) = 51.25, ****P<0.0001); main effect of treatment (F (3, 18) = 208.2, ****P<0.0001). Tukey’s multiple comparison indicated on graph: *p=0.0425 WT Bic vs. WT Veh; ****p<0.0001 WT Veh vs. WT TTX; ****p<0.0001 WT Veh vs. WT CNQX; ****p<0.0001 *Ppt1^-/-^* Veh vs. *Ppt1^-/-^* TTX; ****p<0.0001 *Ppt1^-/-^* Veh vs. *Ppt1^-/-^* CNQX; ****p<0.0001 WT TTX vs. *Ppt1^-/-^* TTX. ***p=0.0004 WT CNQX vs. *Ppt1^-/-^* CNQX. Data represent mean ± SEM.

We first performed a surface biotinylation assay to examine the surface expression of GluA1 after scaling down (Bic), up (TTX), or vehicle control treatment (Veh). As demonstrated in previous studies (Diering et al., 2014; Shepherd et al., 2006), bicuculline treatment of WT cells decreased surface expression of GluA1. However, there was no effect of bicuculline in *Ppt1^-/-^* neurons (**Figure 3A-B**). Interestingly, while TTX treatment induced synaptic upscaling in both WT and *Ppt1^-/-^* neurons, it was exaggerated in *Ppt1^-/-^* cells, as demonstrated by significantly greater surface levels of GluA1 (**Figure 3A-B**). Immunoblots of inputs from the same cell lysates showed an increase in total GluA1 levels to an equal degree in both WT and *Ppt1^-/-^* neurons during upscaling (**Figure 3C-D**). However, we did not observe a significant increase in the total levels of GluA2 in either WT or *Ppt1^-/-^* under these conditions (**Figure 3E**). Collectively, these data indicate that a posttranslational or trafficking mechanism is responsible for the exaggerated GluA1 surface expression.

Next, we employed the same scaling assay and visualized GluA1 surface expression using fluorescent immunolabeling under a non-permeabilizing condition where the antibody does not penetrate the membrane and therefore only surface receptors are labeled (**Figure 3F**). Again, bicuculline treatment did not induce a statistically detectable downscaling of GluA1 puncta (puncta count or percent area/20μm of dendrite) in *Ppt1^-/-^* neurons but reduced surface expression of the GluA1 subunit in WT cells (**Figure 3G, H**). Similar to our biochemical findings (**Figure 3A-B**), TTX treatment induced upscaling in both WT and *Ppt1^-/-^* cells, but the increase was significantly greater in *Ppt1^-/-^* neurons, as we observed by increased GluA1 puncta count and percent area covered along the dendrite (**Figure 3G-H**). These data indicate that *Ppt1^-/-^* neurons *in vitro* exhibit exaggerated synaptic upscaling.

### CP-AMPARs Are Responsible for Exaggerated Upscaling in *Ppt1^-/-^* Neurons

Previous studies showed that both DR and *in vitro* synaptic upscaling procedures increased the postsynaptic GluA1 content and inward rectification of AMPAR-dependent currents in visual cortical neurons, hallmarks of CP-AMPARs (Desai et al., 2002b; Goel and Lee, 2007a; Goel et al., 2006a, 2011). Measurement of GluA1 levels alone does not exclude the possibility that the exaggerated upscaling in *Ppt1^-/-^* neurons is constituted by calcium-impermeable AMPARs. Therefore, we next directly discerned whether CP-AMPARs comprised the synaptic receptor population in overly upscaled *Ppt1^-/-^* synapses. To this end, we performed calcium imaging in WT and *Ppt1^-/-^* primary cortical neurons. We transfected WT and *Ppt1^-/-^* cells with GCaMP3 and exposed them to TTX or vehicle for 48h, then performed a live-cell calcium imaging assay while perfusing the CP-AMPAR specific blocker NASPM (1 or 10μM, where indicated) into the bath (**Figure 4A, Supplementary videos 1-4**). A lab personnel blinded to the study manually selected all active synapses in each video and extracted their ΔF/F_0_ traces (**Figure 4B**). First, corroborating our earlier analyses of GluA1 levels *in vivo* and surface expression *in vitro* (**Figures 2-3**), we did not see a difference in the number of CP-AMPAR transients between WT and *Ppt1^-/-^* vehicle-treated neurons under basal conditions (**Figure 4C, D**). However, while TTX treatment increased the number of synaptic calcium transients in both WT and *Ppt1^-/-^* cells, scaled up *Ppt1^-/-^* neurons exhibited the highest number of baseline calcium transients on average (**Figure 4C**). Importantly, while the proportion of NASPM-sensitive synapses was also similar with vehicle treatment between WT and *Ppt1^-/-^* neurons, upscaled *Ppt1^-/-^* neurons demonstrated significantly more NASPM-sensitive synapses, at both 1μM and 10μM doses, than all other groups (**Figure D, Videos 1-4**). These data indicate that *Ppt1^-/-^* neurons upregulated CP-AMPARs to a greater extent following synaptic upscaling compared to WT cells. Further, the data demonstrate that the activity-dependent trafficking of CP-AMPARs is dysregulated in *Ppt1^-/-^* neurons.

**Figure 4.**
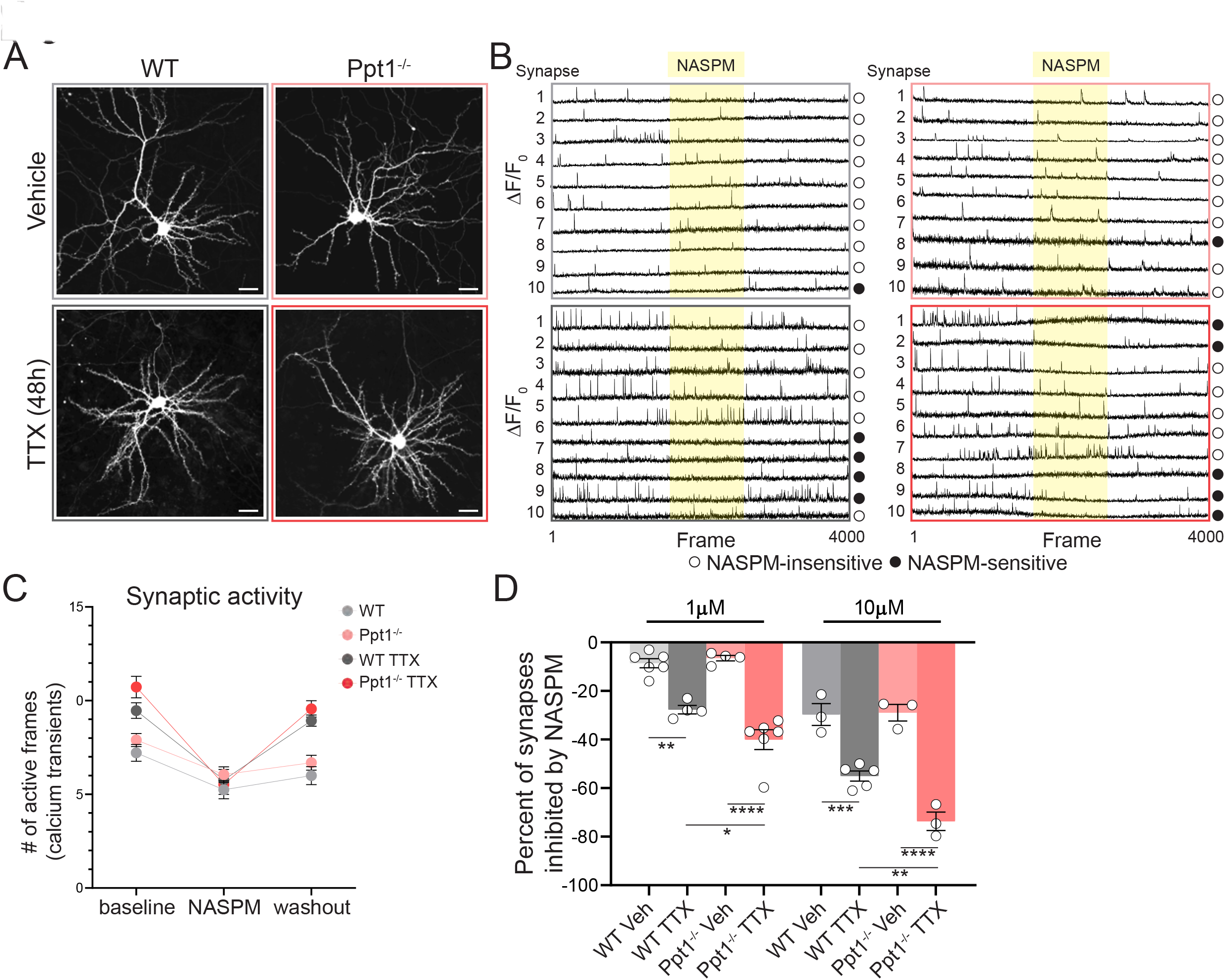
Exaggerated synaptic scaling in *Ppt1^-/-^* neurons is mediated by incorporation of CP-AMPARs. **(A)** Representative calcium imaging (GCaMP) video still frames of vehicle control or TTX-treated (1μm, 48h) WT and *Ppt1^-/-^* cortical neurons. Scale = 20μm. **(B)** Representative ΔF/F_0_ traces for ten synapses in each condition (WT, *Ppt1^-/-^,* TTX-WT, and TTX-*Ppt1^-/-^*). Open circles denote NASPM-insensitive synapses and closed circles denote NASPM-sensitive synapses. **(C)** Quantification of the average synaptic activity (number of frames >3SDs from baseline) for all synapses in one representative cell from each condition prior to, during, and following NASPM (1μM) bath infusion. **(D)** Quantification of the proportion of NASPM-sensitive synapses in WT, TTX-treated WT, *Ppt1^-/-^*, and TTX-treated *Ppt1^-/-^* cortical neurons expressed as the percentage of synapses inhibited during NASPM infusion. N=3-5 cells/group, cells taken from at least 2 independent cultures. One-way ANOVA: F (7, 26) = 52.12, ****P<0.0001. Tukey’s multiple comparison indicated on graph: **p=0.0011 WT-1μm vs. WT TTX-1μm; ****p<0.0001 *Ppt1^-/-^*-1μm vs. *Ppt1^-/-^* TTX-1μm; *p<0.0485 WT TTX-1μm vs. *Ppt1^-/-^*-1μm vs. *Ppt1^-/-^* TTX-1μm; ****p<0.0001 WT-10μm vs. WT TTX-10μm; ****p<0.0001 *Ppt1^-/-^*-10μm vs. *Ppt1^-/-^* TTX- 10μm; **p<0.0061 WT TTX-10μm vs. *Ppt1^-/-^* TTX-10μm. Data represent mean ± SEM.

### Exaggerated Synaptic Upscaling in *Ppt1^-/-^* Neurons Correlate with Impaired Membrane Mobility and an Overload of Palmitoylated GluA1

To examine potential mechanisms by which loss of Ppt1 disrupts GluA1 trafficking during upscaling, we next measured whether AMPAR subunit mobility at the synaptic membrane is impaired using fluorescence recovery after photobleaching (FRAP) analysis of the superecliptic (SEP) GluA1 subunit in WT and *Ppt1^-/-^* neurons (**Figure 5A**). Photobleaching of multiple individual dendritic spines per neuron revealed a significantly slower recovery of photobleached SEP-GluA1 signal at individual synapses in *Ppt1^-/-^* cells (**Figure 5B-C**), resulting in an increased immobile fraction of GluA1 in *Ppt1^-/-^* neurons (**Figure 5D**). Thus, the amplified upscaling is induced partly by a reduced surface mobility of GluA1 within the synaptic membrane of *Ppt1^-/-^* neurons. These findings agree with our hypothesis that loss of Ppt1 function disrupts proteostasis of palmitoylated proteins.

**Figure 5.**
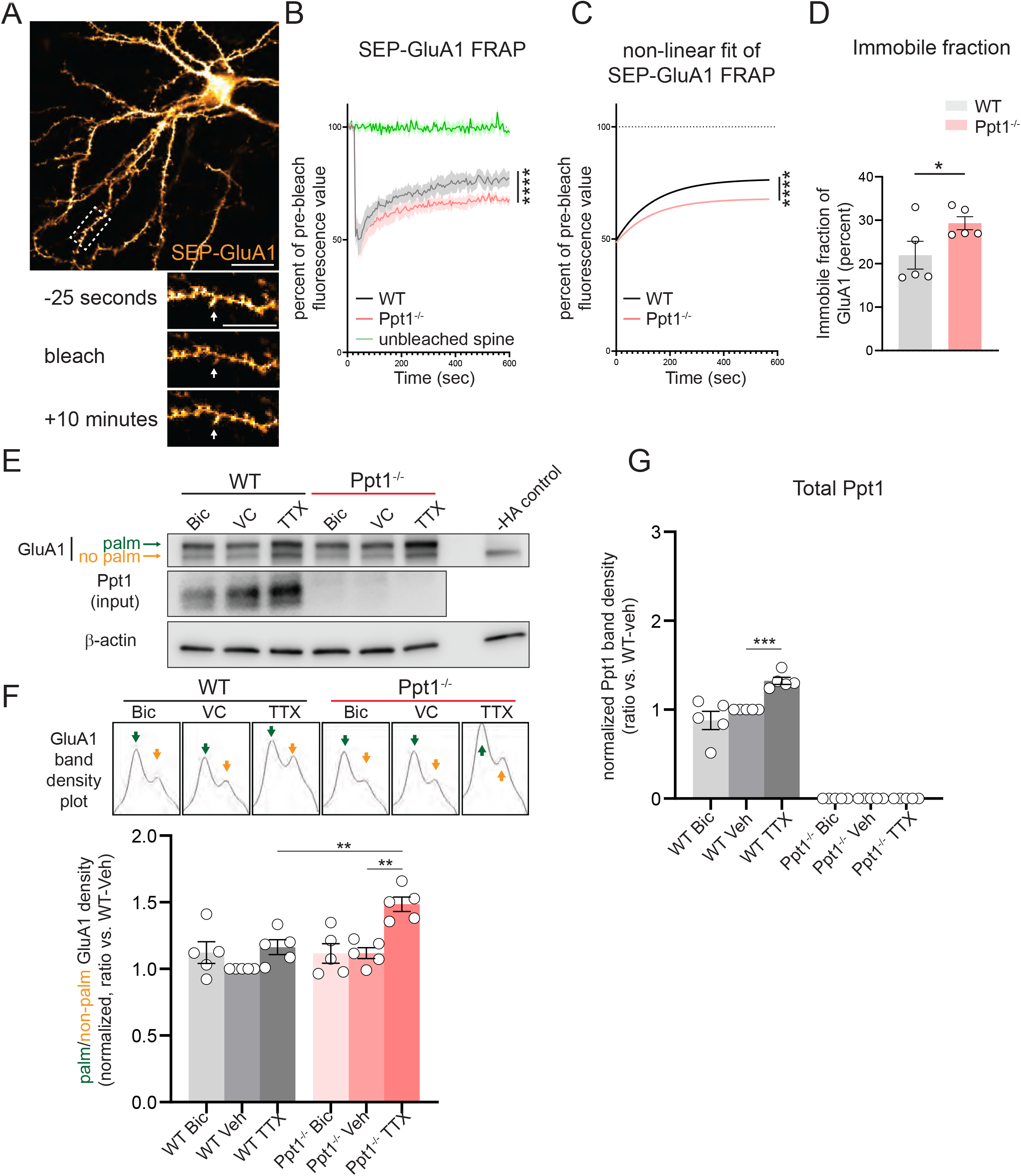
Exaggerated synaptic scaling in *Ppt1^-/-^* cortical neurons is associated with impaired membrane mobility and over-palmitoylation of GluA1. **(A)** Representative image of a SEP-GluA1 transfected neuron (top) and a 20μm segment of dendrite before, just after, and 10min after bleaching of an individual spine (arrow, bottom). Scale=20μm for low magnification images (top), 10μm for high magnification images (bottom). **(B)** Time-dependent recovery of SEP-GluA1 fluorescence after photobleaching (FRAP) in WT and *Ppt1^-/-^* primary cortical neurons. Five-six spines were bleached for each cell and averaged for to make one n. Trace represents average of n=5 cells across 2 independent cultures. **(C)** Non-linear fit of (single phase decay) of SEP-GluA1 fluorescence after photobleaching (FRAP) in WT and *Ppt1^-/-^* primary cortical neurons. Comparison of non-linear fits: F (3, 220) = 542.9, **** p<0.0001. n=5 cells across 2 independent cultures. **(D)** Quantification of the immobile fraction (percent unrecovered compared to pre-bleach fluorescence) of SEP-GluA1 fluorescence after photobleaching (FRAP) for each cell. t-test: *p=0.0356. N=5 cells across 2 independent cultures, 5-6 spines/cell. **(E)** Representative immunoblot of APEGS assay-processed cortical neurons lysates probing for GluA1, GluA2, and β-actin. Representative blot for Ppt1 (input) also shown. Neurons were treated with bicuculline (20μM), vehicle control (DMSO, final concentration 0.1%), or TTX (1μM) for 48h prior to APEGS assay. **(F)** (Top) Representative immunoblot band density plots displaying distinct peaks for the palmitoylated (green arrow) and non-palmitoylated fractions (orange arrow), which were quantified separately and expressed a ratio of palmitoylated/non-palmitoylated GluA1. (Bottom) Quantification of the ratio of palmitoylated/non-palmitoylated GluA1 levels, normalized to β-actin loading control. Two-way ANOVA: interaction genotype x treatment (F (2, 24) = 4.211, *P=0.0271); main effect of genotype (F (1, 24) = 9.559, **P=0.0050); main effect of treatment (F (2, 24) = 11.80, ***P=0.0003). Tukey’s multiple comparison indicated on graph: **p=0.0017 *Ppt1^-/-^* Veh vs. *Ppt1^-/-^* TTX; **p=0.0065 WT TTX vs. *Ppt1^-/-^* TTX. N=5 independent cultures. Data represent mean ± SEM. **(G)** Quantification of the Ppt1 levels, normalized to β-actin loading control. Two-way ANOVA: interaction genotype x treatment (F (2, 24) = 13.07, ***P=0.0001); main effect of genotype (F (1, 24) = 847.4, ****P<0.0001); main effect of treatment (F (2, 24) = 13.07, ***P=0.0001). Tukey’s multiple comparison indicated on graph: ***p=0.0004 WT Veh vs. WT TTX. N=5 independent cultures. Data represent mean ± SEM.

A quantitative palmitoyl-proteomic study showed that the AMPAR subunit GluA1 is a substrate of Ppt1 (Gorenberg et al., 2020) and we show its membrane mobility is affected in *Ppt1^-/-^* neurons. To determine whether GluA1 palmitoylation is accountable for the enhanced incorporation or activity of CP-AMPARs in *Ppt1^-/-^* neurons during synaptic scaling, we performed the Acyl-PEGyl Exchange Gel-Shift (APEGS) assay (Koster et al., 2019; Yokoi et al., 2016) in WT and *Ppt1^-/-^* neurons following scaling down (bicuculline), scaling up (TTX), or vehicle control (**Figure 5E, F**). We measured the density of palmitoylated GluA1 (upper band in immunoblot) to the non-palmitoylated fraction (lower band in immunoblot) to calculate the palmitoylated ratio, for which the density plots are displayed (**Figure 5F**, top). While we did not detect any significant change in GluA1 palmitoylation in WT cells following scaling, similar to a previous report (Yang et al., 2009), we observed a significant increase in the palmitoylated GluA1 ratio in upscaled *Ppt1^-/-^* neurons (**Figure 5F**, bottom). Interestingly, Ppt1 levels exhibit a scaling-dependent effect in WT cells—Ppt1 levels are increased by upscaling (TTX) (**Figure 5E, G**), suggesting that activity-dependent expression of Ppt1 during synaptic scaling in WT neurons prevents GluA1 subunits from uncontrolled palmitoylation and accumulation at the synapse. Further, the exaggerated increase in CP-AMPARs in *Ppt1^-/-^* cortical neurons is likely attributable to aberrant GluA1 palmitoylation that occurs with loss of normal Ppt1 function. However, given the distinct effects of GluA1 palmitoylation at each of its currently recognized three sites on AMPA receptor trafficking, further study is required to determine the specific role of increased palmitoylation on GluA1 during scaling in *Ppt1^-/-^* cells.

### Synaptic Upscaling of GluA1 Is Exaggerated in *Ppt1^-/-^* Neurons *in vivo*

Recognizing that loss of *Ppt1^-/-^* amplified synaptic upscaling *in vitro,* we next assessed levels of GluA1 in LR- WT, LR-*Ppt1^-/-^*, DR-WT, *and* DR-*Ppt1^-/-^* visual cortex to determine if this mechanism contributed to the worsening disease symptoms in DR-*Ppt1^-/-^* animals. Comparing GluA1 levels in DR visual cortices and their typically reared counterparts revealed an increase in the GluA1 content in synaptosomes in DR-WT mice, corroborating previous studies (Goel et al., 2006a, 2011). Importantly, the increase in GluA1 in DR-*Ppt1^-/-^* visual cortices exceeded that of WT animals (**Figure 6A**).

**Figure 6.**
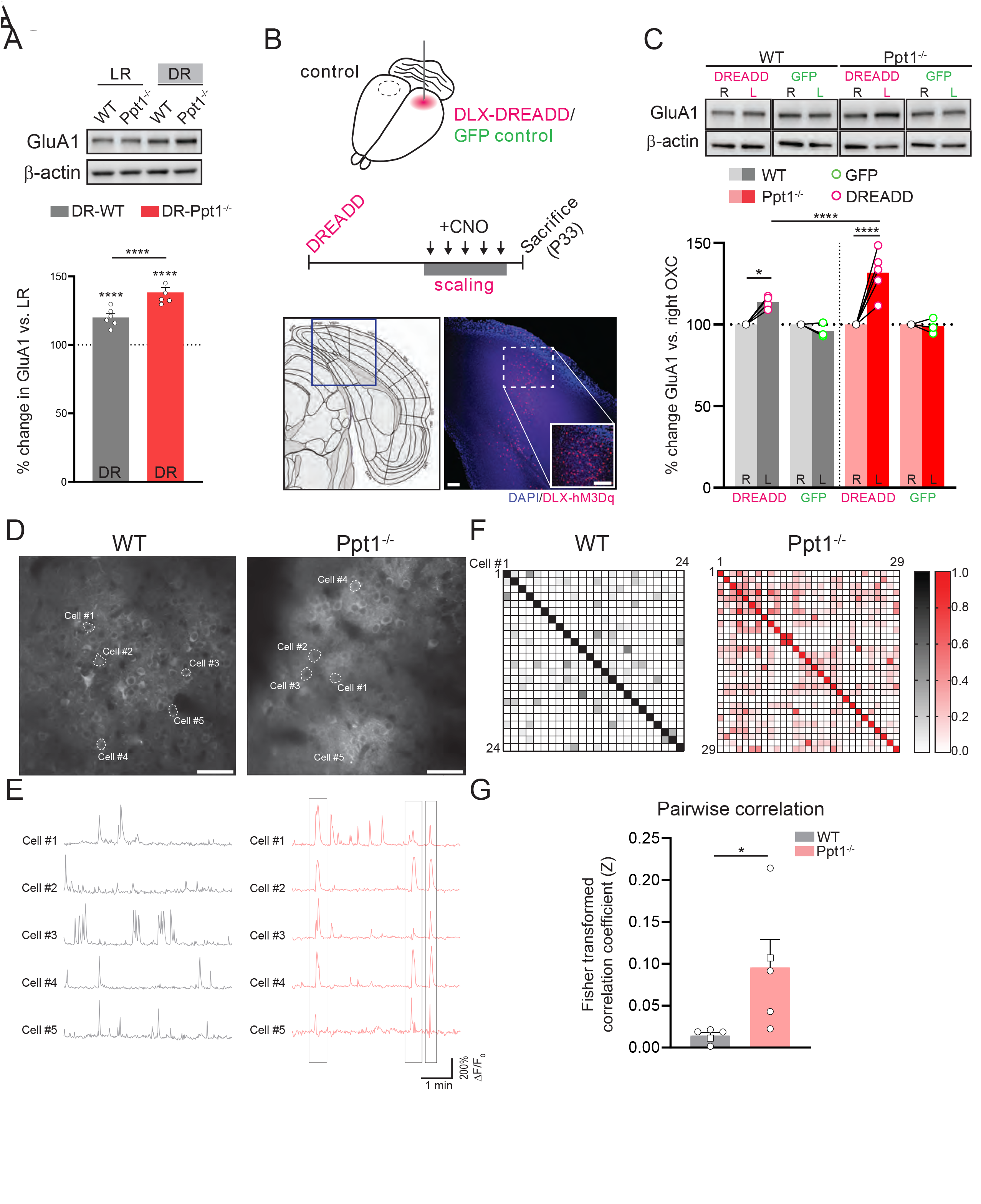
Synaptic upscaling is exaggerated in *Ppt1^-/-^* neurons *in vivo* and leads to hypersynchrony of the visual cortex. **(A)** Representative immunoblot and quantification of GluA1 levels in LR-WT, LR-*Ppt1^-/-^*, DR-WT, and DR- *Ppt1^-/-^* visual cortical synaptosomes. N=6 animals/group. Two-way ANOVA: interaction genotype x rearing condition (F (1, 20) = 17.00, *** P=0.0005); main effect of genotype (F (1, 20) = 17.00, *** P=0.0005); main effect of rearing condition (F (1, 20) = 172.6, P<0.0001). Tukey’s multiple comparison indicated on graph: ****p<0.0001 LR-WT vs. DR-WT; ****p<0.0001 LR-*Ppt1^-/-^* vs. DR-*Ppt1^-/-^*; ****p<0.0001 DR-WT vs. DR-*Ppt1^-/-^*. Data represent mean ± SEM. **(B)** Schematic of the DREADD-induced scaling paradigm and timeline (top). Representative images of DLX- hM3Dq injection in the left visual cortex (bottom right) of a WT mouse aligned to the Allen Brain atlas (bottom left). Scale=50μm for both low and high magnification images. **(C)** Representative immunoblot of synaptosomal GluA1 from WT, WT-DREADD, *Ppt1^-/-^*, and *Ppt1^-/-^*-DREADD occipital cortices and GFP-injected controls (top, all samples from same gel rearranged to match quantification) and quantification (bottom) with. N=3-6 animals/group. Two-way ANOVA: interaction genotype x DREADD (F (1, 17) = 8.740, ** P=0.0088); main effect of genotype (F (1, 17) = 8.740, ** P=0.0088); main effect of DREADD (F (1, 17) = 56.65, **** P<0.0001). Tukey’s multiple comparison indicated on graph: *p= 0.0280 WT vs. DREADD-WT; ****p<0.0001 *Ppt1^-/-^* vs. DREADD-*Ppt1^-/-^*; **p= 0.0057 DREADD-WT vs. DREADD-*Ppt1^-/-^*. **(D)** Representative calcium imaging video still frames of layer 2/3 cortical neuronal somata from a WT and *Ppt1^-/-^* mouse. Scale bar = 50μm. **(E)** Representative ΔF/F_0_ traces for selected cells in each video. Note the coactivity of multiple cells in the *Ppt1^-/-^* brain demarked by grey boxes. **(F)** Correlation matrices of the deconvolved coactivity (Persons r correlation values) of all cells in in the representative videos. Each square represents 1 cell, as denoted in the top left. **(G)** Averaged pair-wise correlation value (Fisher transformed) for all videos, from all animals. Each point represents the average for one mouse. t-test: *p=0.0419. Square points denote the representative correlation matrices from (F). n=5 mice/group

To expand on these findings, we turned to a chemogenetic approach to more precisely induce synaptic upscaling in the visual cortex. We unilaterally activated GABAergic cells in the visual cortex of WT and *Ppt1^-/-^* mice with an excitatory designer receptor exclusively activated by designer drugs (DREADD) under the control of the Dlx5 promoter (Dimidschstein et al., 2017) and administered clozapine-N-oxide (CNO) for five consecutive days (a similar approach to (Wen and Turrigiano, 2021). This method allowed us to chronically suppress local excitatory activity and thereby induce synaptic local synaptic upscaling (**Figure 6B**). We demonstrated, mimicking our results in DR mice, that the DREADD-induced upscaling of synaptosomal GluA1 levels was exaggerated in the *Ppt1^-/-^* mouse visual cortices as compared to WT and GFP-injected *Ppt1^-/-^* controls (**Figure 6C**). Taken together, these data show that synaptic upscaling is amplified in *Ppt1^-/-^* neurons both *in vitro* and *in vivo*.

### *In vivo* Two-Photon Calcium Imaging Reveals Increased Cortical Synchrony in *Ppt1^-/-^* Visual Cortex

Computational studies show that unrestrained homeostatic plasticity causes burst discharges reminiscent of epileptiform activity, a prominent feature of CLN1, and hypersynchrony of artificial cortical circuits (Fröhlich et al., 2008; Houweling et al., 2005). To determine the circuit-level effects of dysregulated synaptic scaling in *Ppt1^-/-^* mice, we employed *in vivo* two-photon calcium imaging in awake mice. We transfected layer 2/3 visual cortical neurons with the calcium indicator, GCaMP6f and fluorescent cell marker, mCherry, in neonates (see Key Resources Table) implanted a cranial window over the left V1 area of injected animals between P21-27, and imaged visual cortical calcium activity at P28-P35 (recovery period from surgery was a minimum of 7 days).

We monitored visual cortical neuron activity for 5-minute epochs in WT and *Ppt1^-/-^* mice and extracted the ΔF/F_0_ trace for each active neuron in each video (**Supplementary videos 5-6**; **Figure 6D, E**). Spontaneous activity in WT and *Ppt1^-/-^* visual cortical neurons showed no significant difference in the total number of active neurons or the average activity of individual neurons per field of view; however, analyzing the pair wise co-activity of all neurons in the videos revealed that compared to WT, the calcium activity of *Ppt1^-/-^* visual cortical neurons demonstrated a significantly increased correlation (Fisher corrected Pearson’s r) (**Figure 6F, G**). These data indicate that although the number of AMPAR-containing excitatory synapses are decreased in *Ppt1^-/-^* mice at P28, the preserved connectivity and neuronal activity drive hypersynchrony of spontaneous activity in the *Ppt1^-/-^* visual cortex.

### Palmitome Study Points to Over-Palmitoylation of Akap5 in *Ppt1^-/-^* Visual Cortex

To further understand the molecular pathways that underlie dysregulated synaptic upscaling in *Ppt1^-/-^* neurons, we employed palmitoyl-proteomics using the acyl-biotin exchange method (Drisdel and Green, 2004; Roth et al., 2006) to visual cortical lysates and synaptosomes (Wan et al., 2007). This analysis detected 512 palmitoylated proteins in lysates and 596 in synaptosomes from WT and *Ppt1^-/-^* visual cortices (**Figure 7A-D**). In lysates, we did not detect any significant changes in the abundance ratio (*Ppt1^-/-^*/WT) for any proteins (**Figure 7A**). In synaptosomes, we detected significantly increased palmitoylation levels of only two proteins, acid ceramidase and cathepsin D, demonstrating that these proteins are consistently overrepresented in the CLN1 brain across several studies (**Figure 7B**) (Atiskova et al., 2019; Chandra et al., 2015; Gorenberg et al., 2020; Sleat et al., 2017). Despite only a few proteins showing statistically significant excessive palmitoylation, we found that most of the identified proteins showed modest overpalmitoylation in the *Ppt1^-/-^* brain, particularly in synaptosomes. Specifically, in *Ppt1^-/-^* lysates, 144 proteins (28.1%) show a >1.2-fold increase in the raw abundance ratio compared to WT, vs 13 (2.5%) showing a <0.80-fold change (**Figure 7C**). The trend is more robust in *Ppt1^-/-^* synaptosomes, with 380 proteins (63.8%) demonstrating a >1.2-fold change increase, while only 31 proteins (5.2%) show a <0.80-fold change reduction (**Figure 7D**). While individual synaptic proteins do not demonstrate statistically significant changes in their palmitoylation by ∼1.5 months, these data implied a trend toward a bulk increase of the palmitoylation level of synaptic proteins with loss of Ppt1 function.

**Figure 7.**
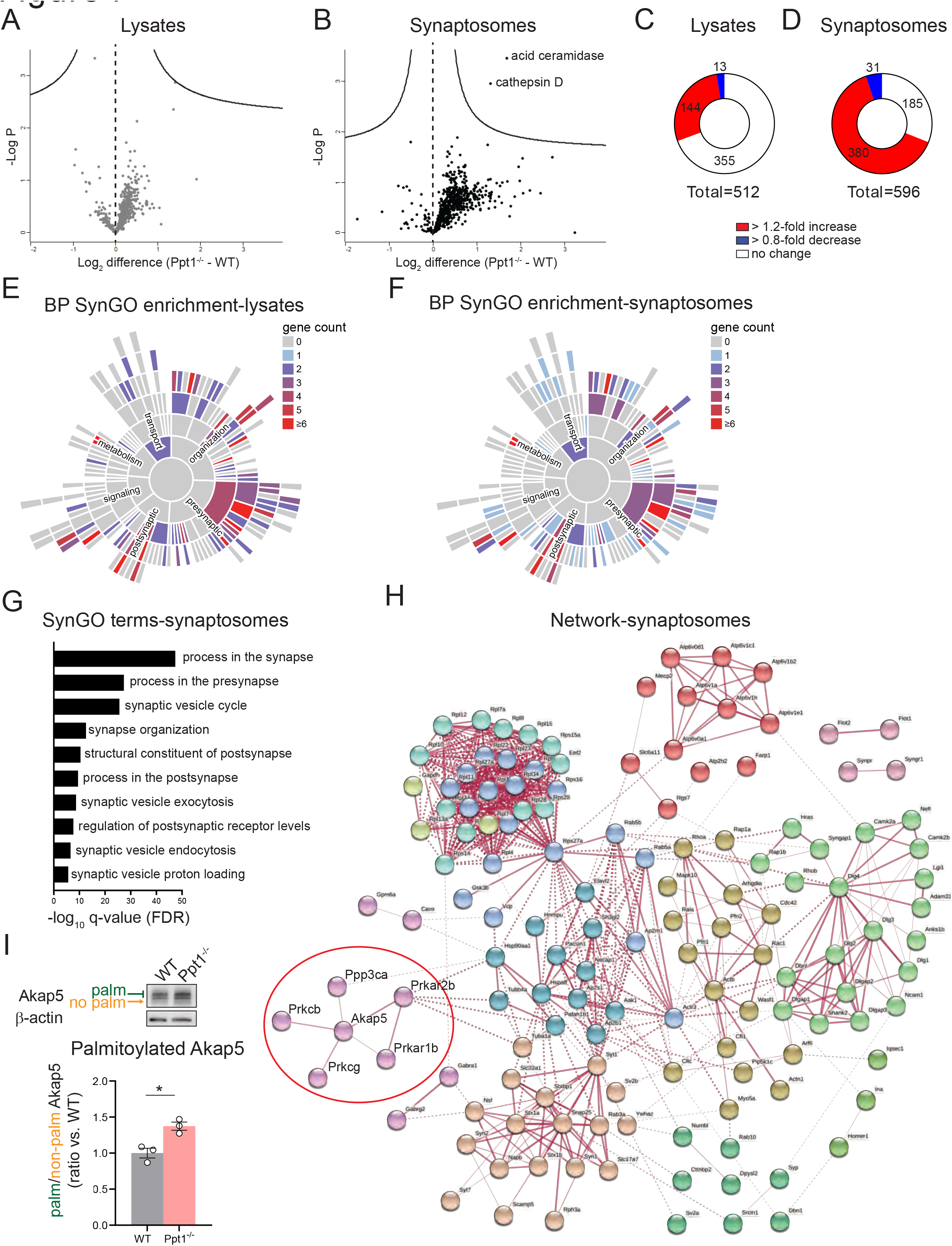
Palmitoyl-proteomics points to excessive palmitoylation of Akap5 and associated signaling proteins in the *Ppt1^-/-^* visual cortex. **(A)** Volcano plot showing the log2 fold change in palmitoyl-protein expression from lysates of WT and *Ppt1^-/-^* visual cortex. N=6 visual cortices/group. **(B)** Volcano plot showing the log2 fold change in palmitoyl-protein expression from synaptosomes of WT and *Ppt1^-/-^* visual cortex. N=6 visual cortices/group. **(C)** Breakdown of the proportion of proteins exhibiting a 1.2-fold (red), 0.8-fold (blue), or no change (white) in the abundance ratio *Ppt1^-/-^*/WT from visual cortical lysates. **(D)** Breakdown of the proportion of proteins exhibiting a 1.2-fold (red), 0.8-fold (blue), or no change (white) in the abundance ratio *Ppt1^-/-^*/WT from visual cortical synaptosomes. **(E)** SynGO annotation of the palmitoyl-proteome of visual cortical lysates from WT and *Ppt1^-/-^* mice. **(F)** SynGO annotation of the palmitoyl-proteome of visual cortical synaptosomes from WT and *Ppt1^-/-^* mice. **(G)** Top ten enriched SynGO terms from proteins increased 1.2-fold in *Ppt1^-/-^* visual cortical synaptosomes. **(H)** Network analysis of the genes increased in *Ppt1^-/-^* synaptosomes by 1.2-fold that were annotated with the top biological process SynGO term “process at the synapse.” Red circle denotes Akap5 pathway. **(I)** Representative immunoblot and quantification of APEGS-processed synaptosomes probing for Akap5 from WT and *Ppt1^-/-^* visual cortices. t-test: *p=0.0120. N=3 mice/group.

To identify the features of synaptic proteins that are overrepresented in the palmitoyl fraction of *Ppt1^-/-^* visual cortices, we input our gene lists for lysates and synaptosomes separately into the online synaptic protein ontology database, SynGO (Koopmans et al., 2019). For both datasets, SynGO analysis demonstrated a robust enrichment for synaptic proteins compared to a whole-brain proteomic background dataset (**Figure 7E, F**), emphasizing the role for palmitoylation at the synapse. The top ten enriched SynGO terms demonstrated substantial overlap between lysates and synaptosomes, which is expected given the enrichment for palmitoylated proteins in both populations. “Process in the synapse,” “process in the presynapse,” “structural constituent of the postsynapse,” “process in the postsynapse,” and “regulation of postsynaptic receptor levels” are represented in the top ten enriched terms in both lysates and synaptosomes (**Figure 7G, S1A**).

To highlight specific pathways that are dysregulated by lack of Ppt1, we performed a network analysis on all genes annotated with the top SynGO term “process in the synapse” using the online protein-protein interaction database, STRING (https://string-db.org). As expected from the SynGO analysis, the networks in lysates and synaptosomes demonstrated substantial overlap (**Figure 7H, S1B**). We noticed a particular cluster that appeared with minor variation in both lysates and synaptosomes, consisting of *Akap5*, the cAMP-dependent protein kinase subunits *Prkacb*, *Prkar1b*, and *Prkar2b*, the protein kinase C subunits *Prkcb*, and *Prkcg*, and the calcineurin subunit *Ppp3ca* (**Figure 7H**, red circle).

Remarkably, the *Akap5* gene encodes the A-kinase anchoring protein 5 (Akap5), a postsynaptic scaffolding protein that anchors protein kinase A, protein kinase C, and calcineurin to the postsynaptic density via its interactions with MAGUK scaffolding proteins such as Sap97, or PSD-95 (Colledge et al., 2000; Robertson et al., 2009), which were also detected in our analysis but showed no change in palmitoylation level, which is consistent with our previous observations (Koster et al., 2019) and the finding that PSD-95 is not a Ppt1 substrate (Yokoi et al., 2016). Due to its close interaction with glutamate receptors, Akap5 creates a signaling microdomain that links neurotransmitter receptor activity to signaling cascades, translating neuronal activity to long-lasting changes in the postsynaptic neuron (Sanderson and Dell’Acqua, 2011). Critically, Akap5 regulates CP-AMPAR incorporation during synaptic scaling (Sanderson et al., 2018). Furthermore, the palmitoylation state of Akap5 is specifically implicated in the regulation of CP-AMPARs during LTP and is induced by seizure *in vivo* (Keith et al., 2012; Purkey et al., 2018; Sanderson et al., 2018). We validated that Akap5 is indeed excessively palmitoylated in the *Ppt1^-/-^* mice by APEGS assay in visual cortical synaptosomes (**Figure 7I**), although we did not detect changes in the total level of Akap5 in lysates or synaptosomes (**Figure S2A, B**). Adding to our previous *in vitro* findings (Koster et al., 2019), we observe an increase in GluN2B palmitoylation in the *Ppt1^-/-^* visual cortex (**Figure S2C**). In contrast, there was no significant increase in GluA1 or GluA2 palmitoylation in *Ppt1^-/-^* animals at this age (**Figure S2C**).

### Nuclear Factor of Activated T-cells (NFAT) Nuclear Translocation Is Increased in Upscaled *Ppt1^-/-^* **Neurons**

Akap5, PKA and calcineurin represent a key postsynaptic signaling hub that regulates synaptic scaling of CP- AMPARs (Sanderson et al., 2018). Upon activation of Akap5-anchored calcineurin, it is free to dephosphorylate the downstream transcription factor nuclear factor activated in T cells NFAT (NFATc3)(Li et al., 2012; Murphy et al., 2019; Sanderson and Dell’Acqua, 2011), which influences the transcription of cytokines, including the proinflammatory molecule and inducer of synaptic scaling, tumor necrosis factor α (TNFα)(Canellada et al., 2006; Stellwagen and Malenka, 2006). Further, neuroinflammation is a key feature of CLN1 in humans and mouse models (Anderson et al., 2013; Macauley et al., 2014; Radke et al., 2015). Therefore, we hypothesized that an overload of palmitoylated Akap5 leads to this CP-AMPAR-calcineurin-NFATc3 pathway bring sensitized in upscaled *Ppt1^-/-^* neurons.

To determine if overly palmitoylated Akap5 leads to hyperactivation of the downstream NFAT pathway in *Ppt1^-/-^* neurons, we performed a nuclear translocation assay as described in Murphy et al., (2014), following 48h pretreatment of TTX to induce upscaling. Briefly, WT and *Ppt1^-/-^* neurons were exposed to an isotonic, high K^+^ (50mM) Tyrode’s solution to depolarize the culture and allowed to recover for 15 minutes before fixation and immunolabeling for NFATc3-GFP (**Figure 8A, B**). The ratio of the integrated fluorescence signal of nuclear to somatic NFATc3-GFP was measured as a proxy for the activation of NFATc3 transcription. While high K^+^- induced depolarization caused the nuclear translocation of NFATc3-GFP in both WT and *Ppt1^-/-^* neurons, the nuclear/soma ratio was significantly higher in *Ppt1^-/-^* cells, indicating a greater responsiveness to depolarization (**Figure 8B, C**). To test whether this effect resulted from an increased contribution of CP-AMPARs in *Ppt1^-/-^* neurons, we performed the same assay and treated a subset of neurons with NASPM (10μm) during the depolarization period. We found that NASPM reduced NFATc3 nuclear translocation to a greater degree in *Ppt1^-/-^* cells compared to WT (**Figure 8B, C**).

**Figure 8.**
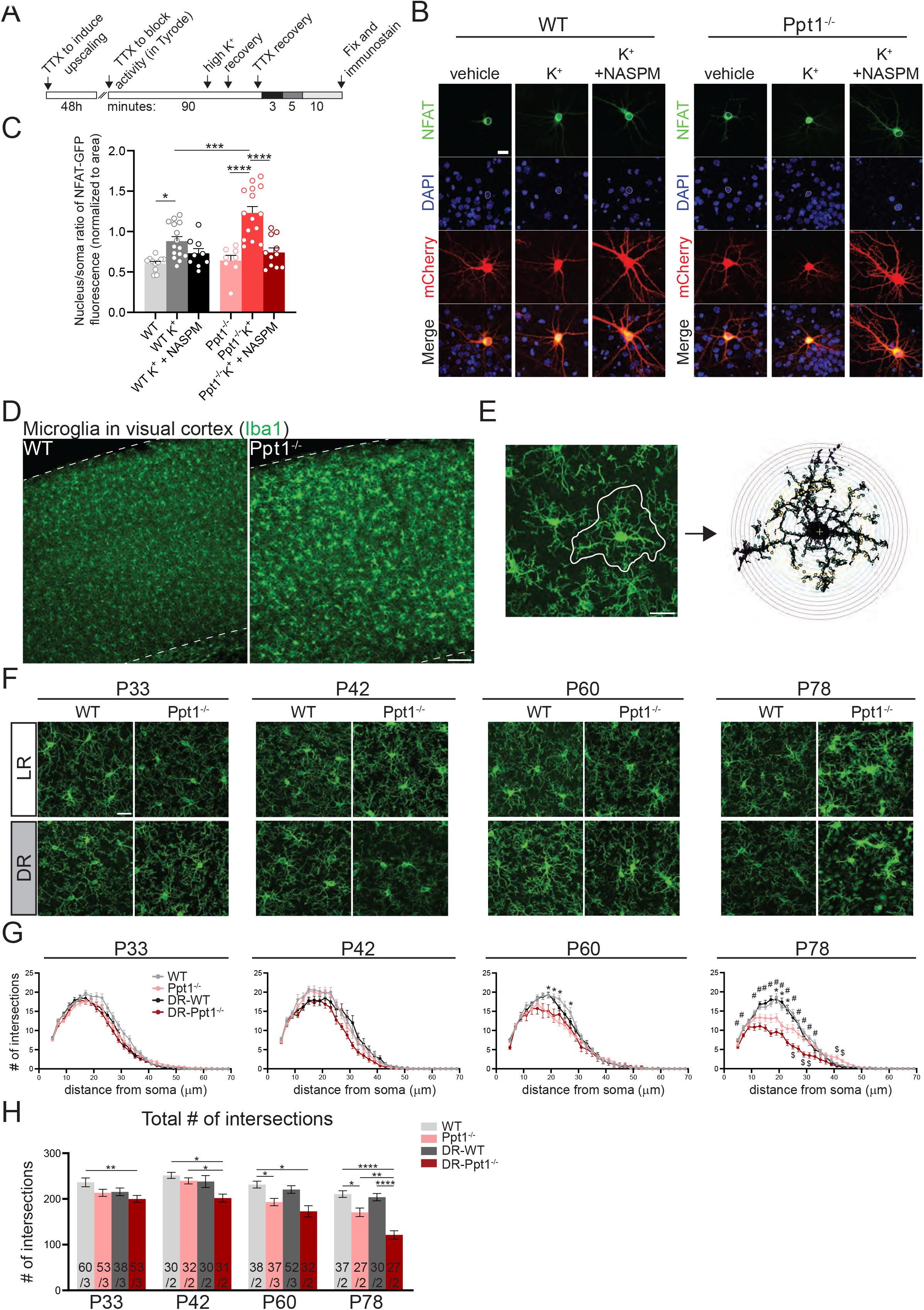
Sensitization of NFAT nuclear translocation in *Ppt1^-/-^* neurons and exacerbated neuroinflammation in dark reared *Ppt1^-/-^* visual cortex. **(A)** Schematic representation of the experimental design for depolarization (K^+^) induction of NFAT activity (nuclear translocation), as previously described (Murphy et al., 2014). **(B)** Representative images of NFAT localization from WT and *Ppt1^-/-^* neurons in response to either sham or K^+^-induced depolarization. Outline of the nucleus (saturated DAPI signal) is drawn in each image. Scale bar= 20μm. **(C)** Quantification of the nucleus/soma ratio of NFAT fluorescence in WT and *Ppt1^-/-^* upscaled neurons in sham depolarization, K^+^ depolarization, and K^+^ depolarization + NASPM (10μM) conditions. Two-way ANOVA: interaction genotype x treatment (F (2, 62) = 4.675, *P=0.0129); main effect of genotype (F (1, 62) = 5.882, *P=0.0182); main effect of treatment (F (2, 62) = 26.23, ****P<0.0001). Tukey’s multiple comparison indicated on graph: *p=0.0245 WT vs. WT K^+^; ****p<0.0001 *Ppt1^-/-^* vs. *Ppt1^-/-^* K^+^; ****p<0.0001 *Ppt1^-/-^* K^+^ vs. *Ppt1^-/-^* K^+^ + NASPM; ***p=0.006 WT K^+^ vs. *Ppt1^-/-^* K^+^. **(D)** Representative low magnification images of Iba1 immunostaining in the visual cortex of WT and *Ppt1^-/-^* visual cortex at P78. Scale bar=100μm **(E)** Representative thresholded microglia image and Sholl analysis overlay. **(F)** Representative high magnification images of Iba1 immunostaining for visual cortical (layer 2/3) microglia in WT, *Ppt1^-/-^,* DR-WT, and DR-*Ppt1^-/-^* mice across age (P28-P78). Scale bar=20μm **(G)** Quantification of microglia branching by Sholl analysis in WT, *Ppt1^-/-^,* DR-WT, and DR-*Ppt1^-/-^* mice across age, where the number of intersections (y-axis) are averaged across cells at the specified distance from the soma (x-axis). Multiple t-tests: *p<0.005 WT vs. *Ppt1^-/-^*; #p<0.005 WT vs. DR-*Ppt1^-/-^*; $p<0.005 *Ppt1^-/-^* vs. DR-*Ppt1^-/-^* (see **Table 1**). N=27-60cells, 2-3 animals per group. Number of cells/animals for each group is listed in panel (H). **(H)** Quantification of the total number of intersections for each group of animals across age. Number of cells/animals in each group are listed on the graph. Two-way ANOVA: **P=0.0017 WT P33 vs. DR-*Ppt1^-/-^* P33; P42 *p=0.0160 WT P42 vs. DR- *Ppt1^-/-^* P42; *p=0.0250 *Ppt1^-/-^* P42 vs. DR- *Ppt1^-/-^* P42; *p=0.0458 WT P60 vs. *Ppt1^-/-^* P60; *p=0.0107 WT P60 vs. DR- *Ppt1^-/-^* P60; *p=0.0197 DR-WT P60 vs. DR- *Ppt1^-/-^* P60; *p=0.0178 WT P78 vs. *Ppt1^-/-^* P78; ****p<0.0001 WT P78 vs. DR- *Ppt1^-/-^* P78; **p=0.0040 *Ppt1^-/-^* P78 vs. DR- *Ppt1^-/-^* P78; ****p<0.0001 DR-WT P78 vs. DR- *Ppt1^-/-^* P78

**Table 1:** Statistics for Sholl measurements in Figure 8G.

| distance<br>from soma<br>(mm) | P33 |  |  |  |  |  |  |  |
| --- | --- | --- | --- | --- | --- | --- | --- | --- |
|  | WT vs. *Ppt1*<sup>-/-</sup> |  | WT vs. DR-WT |  | WT vs. DR-*Ppt1*<sup>-/-</sup> |  | *Ppt1*<sup>-/-</sup> vs. *Ppt1*<sup>-/-</sup> |  |
|  | signif<br>icant<br>? | p-value<br>(FDR<br>corrected) | signif<br>icant<br>? | p-value<br>(FDR<br>corrected) | signif<br>icant<br>? | p-value<br>(FDR<br>corrected) | signif<br>icant<br>? | p-value<br>(FDR<br>corrected) |
| 5 | No | 0.218615 | No | 0.731765 | No | 0.342738 | No | 0.764286 |
| 7 | No | 0.115316 | No | 0.665366 | No | 0.023971 | No | 0.507645 |
| 9 | No | 0.023024 | No | 0.757519 | No | 0.070949 | No | 0.632289 |
| 11 | No | 0.016119 | No | 0.749431 | No | 0.320358 | No | 0.16471 |
| 13 | No | 0.443392 | No | 0.433162 | No | 0.949456 | No | 0.435926 |
| 15 | No | 0.169529 | No | 0.885876 | No | 0.678793 | No | 0.330704 |
| 17 | No | 0.019493 | No | 0.203353 | No | 0.021276 | No | 0.923168 |
| 19 | No | 0.238231 | No | 0.184467 | No | 0.193691 | No | 0.972754 |
| 21 | No | 0.103026 | No | 0.081512 | No | 0.013395 | No | 0.415518 |
| 23 | No | 0.088268 | No | 0.065691 | No | 0.005041 | No | 0.238345 |
| 25 | No | 0.017733 | No | 0.207134 | No | 0.005962 | No | 0.522911 |
| 27 | No | 0.100663 | No | 0.332808 | No | 0.008963 | No | 0.1873 |
| 29 | No | 0.042891 | No | 0.127728 | No | 0.001193 | No | 0.161231 |
| 31 | No | 0.093855 | No | 0.056172 | No | 0.005948 | No | 0.184524 |
| 33 | No | 0.095839 | No | 0.068832 | No | 0.006357 | No | 0.199908 |
| 35 | No | 0.756193 | No | 0.084214 | No | 0.033589 | No | 0.072587 |
| 37 | No | 0.460413 | No | 0.448059 | No | 0.083245 | No | 0.233528 |
| 39 | No | 0.996764 | No | 0.820076 | No | 0.545421 | No | 0.508027 |
| 41 | No | 0.151357 | No | 0.95319 | No | 0.895777 | No | 0.214591 |
| 43 | No | 0.177366 | No | 0.98 | No | 0.47443 | No | 0.042373 |
| 45 | No | 0.174571 | No | 0.374394 | No | 0.757919 | No | 0.101548 |
| 47 | No | 0.220771 | No | 0.307062 | No | 0.59165 | No | 0.069981 |
| 49 | No | 0.394463 | No | 0.218616 | No | 0.981856 | No | 0.378607 |
| 51 | No | 0.070006 | No | 0.428862 | No | 0.319585 | No | 0.453046 |
| 53 | No | 0.067068 | No | 0.969558 | No | 0.512291 | No | 0.175332 |
| **55** | No | 0.037601 | No | 0.34815 | No | 0.126837 | No | 0.411259 |
| **57** | No | 0.060524 | No | 0.210592 | No | 0.289357 | No | 0.10001 |
| **59** | No | 0.056984 | No | 0.210592 | No | n/a | No | 0.073701 |
| **61** | No | 0.085294 | No | 0.210592 | No | n/a | No | 0.105944 |
| **63** | No | 0.152835 | No | 0.210592 | No | n/a | No | 0.179282 |
| **65** | No | 0.17833 | No | n/a | No | n/a | No | 0.206114 |
| **67** | No | 0.289357 | No | n/a | No | n/a | No | 0.319632 |
| **69** | No | 0.289357 | No | n/a | No | n/a | No | 0.319632 |

**P42**
| distance<br>from soma<br>(mm) | WT vs. *Ppt1*<sup>-/-</sup> |  | WT vs. DR-WT |  | WT vs. DR-*Ppt1*<sup>-/-</sup> |  | *Ppt1*<sup>-/-</sup> vs. *Ppt1*<sup>-/-</sup> |  |
| --- | --- | --- | --- | --- | --- | --- | --- | --- |
|  | signif<br>icant<br>? | p-value<br>(FDR<br>corrected) | signif<br>icant<br>? | p-value<br>(FDR<br>corrected) | signif<br>icant<br>? | p-value<br>(FDR<br>corrected) | signif<br>icant<br>? | p-value<br>(FDR<br>corrected) |
| **5** | No | 0.396741 | No | 0.921723 | No | 0.258172 | No | 0.785052 |
| **7** | No | 0.248326 | No | 0.195573 | No | 0.429474 | No | 0.51606 |
| **9** | No | 0.582495 | No | 0.55886 | No | 0.827879 | No | 0.733264 |
| **11** | No | 0.20303 | No | 0.089469 | No | 0.001971 | No | 0.017393 |
| **13** | No | 0.595972 | No | 0.072207 | No | 0.042873 | No | 0.037278 |
| **15** | No | 0.769439 | No | 0.026501 | No | 0.019756 | No | 0.020678 |
| **17** | No | 0.720407 | No | 0.092106 | No | 0.034111 | No | 0.031902 |
| **19** | No | 0.652024 | No | 0.121929 | No | 0.147108 | No | 0.251252 |
| **21** | No | 0.751749 | No | 0.499174 | No | 0.093399 | No | 0.026346 |
| **23** | No | 0.684518 | No | 0.158393 | No | 0.034383 | No | 0.028827 |
| **25** | No | 0.853868 | No | 0.651096 | No | 0.038439 | No | 0.00732 |
| **27** | No | 0.779986 | No | 0.535067 | No | 0.028959 | No | 0.004309 |
| **29** | No | 0.159493 | No | 0.719335 | No | 0.023159 | No | 0.119059 |
| **31** | No | 0.91702 | No | 0.342562 | No | 0.039199 | No | 0.004069 |
| **33** | No | 0.606311 | No | 0.345113 | No | 0.1277 | No | 0.011961 |
| **35** | No | 0.410562 | No | 0.853032 | No | 0.032665 | No | 0.079759 |
| **37** | No | 0.445763 | No | 0.797802 | No | 0.121882 | No | 0.248505 |
| **39** | No | 0.227193 | No | 0.759825 | No | 0.063528 | No | 0.420187 |
| **41** | No | 0.719807 | No | 0.317051 | No | 0.094788 | No | 0.18128 |
| 43 | No | 0.709899 | No | 0.929205 | No | 0.193721 | No | 0.340131 |
| 45 | No | 0.183202 | No | 0.652983 | No | 0.463794 | No | 0.405537 |
| 47 | No | 0.108852 | No | 0.886934 | No | 0.107949 | No | 0.971079 |
| 49 | No | 0.957727 | No | 0.55886 | No | 0.976299 | No | 0.975908 |
| 51 | No | 0.257453 | No | 0.162403 | No | 0.077064 | No | 0.542596 |
| 53 | No | n/a | No | n/a | No | n/a | No | n/a |
| 55 | No | n/a | No | n/a | No | n/a | No | n/a |
| 57 | No | n/a | No | n/a | No | n/a | No | n/a |
| 59 | No | n/a | No | n/a | No | n/a | No | n/a |
| 61 | No | n/a | No | n/a | No | n/a | No | n/a |
| 63 | No | n/a | No | n/a | No | n/a | No | n/a |
| 65 | No | n/a | No | n/a | No | n/a | No | n/a |
| 67 | No | n/a | No | n/a | No | n/a | No | n/a |
| 69 | No | n/a | No | n/a | No | n/a | No | n/a |

**P60**
| distance<br>from soma<br>(mm) | WT vs. *Ppt1*<sup>-/-</sup> |  | WT vs. DR-WT |  | WT vs. DR-*Ppt1*<sup>-/-</sup> |  | *Ppt1*<sup>-/-</sup> vs. *Ppt1*<sup>-/-</sup> |  |
| --- | --- | --- | --- | --- | --- | --- | --- | --- |
|  | signif<br>icant<br>? | p-value<br>(FDR<br>corrected) | signif<br>icant<br>? | p-value<br>(FDR<br>corrected) | signif<br>icant<br>? | p-value<br>(FDR<br>corrected) | signif<br>icant<br>? | p-value<br>(FDR<br>corrected) |
| 5 | No | 0.477723 | No | 0.664789 | No | 0.006412 | No | 0.020372 |
| 7 | No | 0.420831 | No | 0.714423 | No | 0.611685 | No | 0.816119 |
| 9 | No | 0.771676 | No | 0.457596 | No | 0.824366 | No | 0.984798 |
| 11 | No | 0.058344 | No | 0.136095 | No | 0.112691 | No | 0.661245 |
| 13 | No | 0.186948 | No | 0.810744 | No | 0.489697 | No | 0.935345 |
| 15 | No | 0.204972 | No | 0.894806 | No | 0.100379 | No | 0.429375 |
| 17 | Yes | 0.006693 | No | 0.878188 | No | 0.051701 | No | 0.755556 |
| 19 | Yes | 0.000192 | No | 0.96489 | No | 0.017115 | No | 0.745268 |
| 21 | Yes | 0.000057 | No | 0.60339 | No | 0.010933 | No | 0.627345 |
| 23 | Yes | 0.000011 | No | 0.099953 | No | 0.019504 | No | 0.157207 |
| 25 | Yes | 0.006549 | No | 0.219129 | No | 0.013333 | No | 0.71113 |
| 27 | Yes | 0.000707 | No | 0.13088 | No | 0.021889 | No | 0.449459 |
| 29 | No | 0.049382 | No | 0.478572 | No | 0.099788 | No | 0.974565 |
| 31 | No | 0.507138 | No | 0.822665 | No | 0.093195 | No | 0.261626 |
| 33 | No | 0.63427 | No | 0.431995 | No | 0.884664 | No | 0.566253 |
| 35 | No | 0.56112 | No | 0.800801 | No | 0.517757 | No | 0.887647 |
| 37 | No | 0.679379 | No | 0.562184 | No | 0.93761 | No | 0.679088 |
| 39 | No | 0.358608 | No | 0.568232 | No | 0.794186 | No | 0.612062 |
| 41 | No | 0.457378 | No | 0.681405 | No | 0.782115 | No | 0.356352 |
| 43 | No | 0.444392 | No | 0.83137 | No | 0.972842 | No | 0.543739 |
| 45 | No | 0.611317 | No | 0.865527 | No | 0.850617 | No | 0.865063 |
| 47 | No | 0.847097 | No | 0.366599 | No | 0.816893 | No | 0.6132 |
| 49 | No | 0.332585 | No | 0.46502 | No | 0.327636 | No | 0.714768 |
| 51 | No | 0.4539 | No | 0.42715 | No | 0.713331 | No | 0.730071 |
| 53 | No | 0.32822 | No | 0.367752 | No | 0.659737 | No | 0.185837 |
| 55 | No | 0.164896 | No | 0.944479 | No | 0.773024 | No | 0.117212 |
| 57 | No | 0.164896 | No | 0.950238 | No | 0.382432 | n/a | n/a |
| 59 | No | 0.164896 | No | 0.705488 | No | 0.382432 | n/a | n/a |
| 61 | n/a | n/a | No | 0.549339 | n/a | n/a | n/a | n/a |
| 63 | n/a | n/a | n/a | n/a | n/a | n/a | n/a | n/a |
| 65 | n/a | n/a | n/a | n/a | n/a | n/a | n/a | n/a |
| 67 | n/a | n/a | n/a | n/a | n/a | n/a | n/a | n/a |
| 69 | n/a | n/a | n/a | n/a | n/a | n/a | n/a | n/a |

P78
| distance<br>from soma<br>(mm) | WT vs. *Ppt1*<sup>-/-</sup> |  | WT vs. DR-WT |  | WT vs. DR-*Ppt1*<sup>-/-</sup> |  | *Ppt1*<sup>-/-</sup> vs. *Ppt1*<sup>-/-</sup> |  |
| --- | --- | --- | --- | --- | --- | --- | --- | --- |
|  | signif<br>icant<br>? | p-value<br>(FDR<br>corrected) | signif<br>icant<br>? | p-value<br>(FDR<br>corrected) | signif<br>icant<br>? | p-value<br>(FDR<br>corrected) | signif<br>icant<br>? | p-value<br>(FDR<br>corrected) |
| 5 | No | 0.042696 | No | 0.824672 | Yes | 0.001025 | No | 0.397884 |
| 7 | No | 0.963773 | No | 0.972826 | Yes | 0.008919 | No | 0.011975 |
| 9 | No | 0.449968 | No | 0.920051 | No | 0.124699 | No | 0.494187 |
| 11 | No | 0.124403 | No | 0.502538 | Yes | 0.000246 | No | 0.063402 |
| 13 | Yes | 0.010318 | No | 0.369974 | Yes | 0.000008 | No | 0.077599 |
| 15 | Yes | 0.006424 | No | 0.396186 | Yes | <0.000001 | No | 0.031292 |
| 17 | Yes | 0.002359 | No | 0.33852 | Yes | <0.000001 | No | 0.003346 |
| **19** | Yes | 0.000232 | No | 0.584397 | Yes | <0.000001 | No | 0.008191 |
| **21** | Yes | 0.000151 | No | 0.257106 | Yes | <0.000001 | No | 0.095409 |
| **23** | Yes | 0.000182 | No | 0.765976 | Yes | <0.000001 | No | 0.006013 |
| **25** | Yes | 0.006791 | No | 0.424461 | Yes | <0.000001 | Yes | 0.000851 |
| **27** | No | 0.054901 | No | 0.972101 | Yes | 0.000003 | No | 0.003857 |
| **29** | No | 0.177626 | No | 0.294129 | Yes | 0.000003 | Yes | 0.000265 |
| **31** | No | 0.340478 | No | 0.341287 | Yes | 0.000121 | Yes | 0.001447 |
| **33** | No | 0.43063 | No | 0.129298 | Yes | 0.003815 | No | 0.014931 |
| **35** | No | 0.80783 | No | 0.916156 | No | 0.079029 | No | 0.04226 |
| **37** | No | 0.54775 | No | 0.928594 | No | 0.243799 | No | 0.062919 |
| **39** | No | 0.214463 | No | 0.554299 | No | 0.180117 | No | 0.005381 |
| **41** | No | 0.019274 | No | 0.465019 | No | 0.28035 | Yes | 0.000484 |
| **43** | No | 0.028008 | No | 0.470463 | No | 0.384071 | Yes | 0.002221 |
| **45** | No | 0.026521 | No | 0.525234 | No | 0.427498 | No | 0.013471 |
| **47** | No | 0.497711 | No | 0.32936 | No | 0.34659 | No | 0.061376 |
| **49** | No | 0.511864 | No | 0.986315 | No | 0.957238 | No | 0.514134 |
| **51** | No | 0.176284 | No | 0.659492 | No | 0.479857 | No | 0.632634 |
| **53** | No | 0.717842 | No | 0.591459 | No | 0.919034 | No | 0.790325 |
| **55** | No | 0.799579 | No | 0.671725 | No | 0.39728 | No | 0.178828 |
| **57** | No | 0.39728 | No | 0.894499 | No | 0.39728 | n/a | n/a |
| **59** | No | 0.39728 | No | 0.672316 | No | 0.39728 | n/a | n/a |
| **61** | n/a | n/a | No | 0.269998 | n/a | n/a | n/a | n/a |
| **63** | n/a | n/a | n/a | n/a | n/a | n/a | n/a | n/a |
| **65** | n/a | n/a | n/a | n/a | n/a | n/a | n/a | n/a |
| **67** | n/a | n/a | n/a | n/a | n/a | n/a | n/a | n/a |
| **69** | n/a | n/a | n/a | n/a | n/a | n/a | n/a | n/a |

**Table 2:** Statistics for Figure S3C.

| dLGN (P21-P42) |  |  |
| --- | --- | --- |
|  | WT vs. *Ppt1*<sup>-/-</sup> |  |
| distance from soma (μm) | significant? | p-value |
| 5 | No | 0.549601 |
| 7 | No | 0.964156 |
| 9 | No | 0.668587 |
| 11 | No | 0.205016 |
| 13 | No | 0.616089 |
| 15 | No | 0.098918 |
| 17 | No | 0.062513 |
| 19 | No | 0.065219 |
| 21 | No | 0.020446 |
| 23 | No | 0.530833 |
| 25 | No | 0.926505 |
| 27 | Yes | 0.001545 |
| 29 | No | 0.01102 |
| 31 | Yes | 0.000034 |
| 33 | Yes | 0.000659 |
| 35 | No | 0.012683 |
| 37 | No | 0.042333 |
| 39 | No | 0.032263 |
| 41 | No | 0.266395 |
| 43 | No | 0.666866 |
| 45 | No | 0.816703 |
| 47 | No | 0.83697 |
| 49 | No | 0.751296 |
| 51 | No | 0.870362 |
| 53 | No | 0.859203 |
| 55 | No | 0.950965 |
| 57 | No | 0.990564 |
| 59 | No | 0.969813 |
| 61 | No | 0.969813 |
| 63 | No | 0.969813 |
| 65 | No | 0.969813 |
| 67 | No | 0.969813 |
| 69 | No | 0.969813 |

**Table 3.** Key Resources Table

| Table 3. Key Resources Table |  |  |  |  |
| --- | --- | --- | --- | --- |
| Reagent type (species) or resource | Designation | Source or reference | Identifiers | Additional information |
| strain, strain background (*Mus musculus*) | B6;129-Ppt1<sup>tm1Hof</sup>/J | Jax stock #: 004313 | Gupta, PNAS, 2001; RRID: MGI:004313 |  |
| antibody | Mouse monoclonal anti-GluA1 | Millipore Sigma | Cat: MAB2263; RRID: AB_11212678 | 1:500 for biotinylation, 1:1000 for all else |
| antibody | Mouse monoclonal anti-GluA2 | UC Davis/NIH NeuroMab Facility | Cat: 75-002; RRID: AB_2232661 | 1:500 for biotinylation, 1:1000 for all else |
| antibody | Rabbit polyclonal anti-Akap79/150 | Gift from Mark Dell'Acqua | RRID: AB_2532138 | 1:1000 |
| antibody | Mouse monoclonal anti-GluN2B | UC Davis/NIH NeuroMab Facility | Cat: 75/097; RRID: AB_10673405 | (1:1000) |
| antibody | Rabbit polyclonal anti-PPT1 | Gift from Sandra Hofmann |  | 1:500 |
| antibody | Mouse monoclonal anti-β-actin | ThermoFisher Scientific | Cat: A2228; RRID: AB_476697 | 1:2000 |
| antibody | Donkey anti-Rabbit IgG Secondary Antibody, HRP | ThermoFisher Scientific | Cat: SA1-200; RRID: AB_325994 | 1:1000 for biotinylation; 1:5000 or 1:10000 for all else |
| antibody | Rabbit polyclonal anti-MAP2 | Millipore Sigma | Cat: AB5622; RRID: AB_91939 | 1:400 |
| antibody | Peroxidase AffiniPure Goat Anti-Mouse IgG | Jackson ImmunoResearch | Cat: 115-035-146; RRID: AB_2307392 | 1:1000 for biotinylation; 1:5000 or 1:10000 for all else |
| antibody | Rat monoclonal anti-LAMP2 | abcam | Cat: ab13524; RRID: AB_2134736 | 1:400 |
| antibody | GFP Polyclonal Antibody, Alexa Fluor 488 | ThermoFisher Scientific | Cat: A-21311; RRID: AB_221477 | 1:1000 |
| antibody | Rabbit polyclonal anti-Iba1 | Fujifilm/Wako | Cat: 019-1974; RRID: AB_839504 | 1:1000 |
| antibody | Mouse monoclonal anti-transferrin | ThermoFisher Scientific | Cat: A-11130; RRID: AB_2534136) | 1:500 |
| antibody | Gt anti-rabbit 633 | ThermoFisher Scientific | Cat: A-21070; RRID: AB_2535731 | 1:400 |
| antibody | Gt anti-mouse 488 | ThermoFisher Scientific | Cat: A28175;<br>RRID:<br>AB_2536161 | 1:400 |
| Virus | AAV8-hDlx-Gq-DREADD-dTomato-Fishell-4 | Vector Biolabs | Addgene (83897-AAV9) | Packaged in AAV8; Gift from Dr. Kuei Tseng |
| Virus | AAV8-pAAV-mDlx-GFP-Fishell-1 | Vector Biolabs | Addgene (83900-AAV9) | Packaged in AAV8; Gift from Dr. Kuei Tseng |
| Virus | AAV9-pENN.AAV.CamKII.GCaMP6f.WPRE.SV40 | Addgene | Cat: 100834-AAV9 |  |
| commercial assay or kit | Dynabeads™ M-280 Streptavidin | ThermoFisher Scientific | Cat: 11205D |  |
| recombinant DNA reagent | pEF-GFP | Addgene | Plasmid: 11154 | Gift from Drs. Matsuda and Cepko |
| recombinant DNA reagent | GCaMP3 | Addgene | Plasmid: 22692 | Dr. Looger |
| recombinant DNA reagent | GFP-mNFATc3 | Gift from Mark Dell'Acqua |  | Mouse NFATc3 with GFP tag |
| recombinant DNA reagent | CAG-mCherry |  |  |  |
| commercial assay or kit | Dynabeads™ M-280 Streptavidin | ThermoFisher Scientific | Cat: 11205D |  |
| chemical compound, drug | Tetrodotoxin-citrate (TTX) | Tocris | Cat: 1069 | Treatment: 1μM |
| chemical compound, drug | NASPM | Tocris | Cat: 2766 | Treatment: 1μM or 10μM where indicated |
| chemical compound, drug | 5KDa maleimide PEG (for APEGs) | NOF America Corporation | Cat: ME-050MA |  |
| chemical compound, drug | (+)-Bicuculline | Tocris | Cat: 0130 | Treatment: 20μM |
| chemical compound, drug | Cholera toxin subunit B 488 | ThermoFisher Scientific | Cat: C34775 | 1mg/ml in PBS |
| chemical compound, drug | Cholera toxin subunit B 488 | ThermoFisher Scientific | Cat: C34776 | 1mg/ml in PBS |
| chemical compound, drug | Clozapine N-oxide | NIH |  | Gift from Dr. Kuei Tseng |
| software, algorithm | Fiji |  |  |  |
| software, algorithm | Matlab 2020a | Mathworks |  | EZcalcium (Cantu et al., 2020) |
| software, algorithm | Prism 9.0.1 | Graphpad |  |  |

### Dark Rearing Exacerbates Neuroinflammation in the Visual Cortex of *Ppt1^-/-^* Mice

Given NFATc3 activation is enhanced in *Ppt1^-/-^* neurons and induction of synaptic scaling *in vivo* by DR leads to exacerbated disease symptoms in *Ppt1^-/-^* visual cortex, we reasoned that DR would worsen the neuroinflammation characteristic of CLN1 (Kielar et al., 2007; Macauley et al., 2014; Palmer et al., 2013). To test this hypothesis, we stained LR-WT, LR-*Ppt1^-/-^,* DR-WT, and DR-*Ppt1^-/-^* brains for the microglia marker, Iba1 (**Figure 8D**), and performed Sholl analysis (**Figure 8E**) on individual cells in the visual cortex (**Figure 8F**), since a classical sign of inflammatory microglial activation is a loss of ramification and ameboid shape (Gehrmann et al., 1995; Giulian, 1987). At P33 and P42, there were no differences between WT and *Ppt1^-/-^* cells in either the number of intersections as a function of distance from the cell soma (**Figure 8G**) or in terms of the total number of intersections (**Figure 8H**). By P60, however, there was a significant reduction in the number of intersections in *Ppt1^-/-^* mice compared to WT that persisted to P78, suggesting microglial morphology analysis is a sensitive measure of the neuroinflammatory phenotype in CLN1 mice (**Figure 8G, H**).

Though we did not detect significant change in microglial morphology in the visual cortex of *Ppt1^-/-^* mice until P60, it is well-known that gliosis and neurodegeneration follows a systematic pattern in CLN1, affecting the thalamus (especially the visual thalamus) before cortex (Kielar et al., 2007). Therefore, we examined the neuroinflammatory phenotype of microglia in the visual thalamus, the dorsal lateral geniculate nucleus (dLGN) at ∼P33 (**Figure S3A-B**). Microglia in this region are crucial for synaptic refinement during development of the visual circuit (Katz and Shatz, 1996; Shatz, 1990), so we focused our analysis on microglia at the borders of ipsilateral and contralateral retinal projections reaching the dLGN. We found a subtle change in microglia morphology that suggests an emerging neuroinflammatory phenotype in the dLGN of *Ppt1^-/-^* mice (**Figure S3C**), although measuring the total number of intersections did not show a significant difference between WT and *Ppt1^-/-^* mice (**Figure S3D**).

Importantly, as early as P33, DR-*Ppt1^-/-^* microglia exhibited a decrease in the total number of intersections compared to WT cells. At P42, before any neurological deficit was detected in LR-*Ppt1^-/-^* mice, DR-*Ppt1^-/-^* mice exhibited significantly reduced microglial processes compared to LR-WT, DR-WT., Moreover, LR-*Ppt1^-/-^*animals (**Figure 8G, H**), demonstrated an acceleration of the neuroinflammatory phenotype in DR animals. This effect was even more robust at P78 (**Figure 8G, H**). These data demonstrate that DR worsens neuroinflammation in the visual cortex of *Ppt1^-/-^* mice, accounting for the accelerated brain atrophy/neurodegeneration we observed (**Figure 1**).

## Discussion

We demonstrate herein that DR *Ppt1^-/-^* animals exacerbated CLN1 disease pathology (**Figure 1**), contrary to the beneficial effects observed in other mouse model of neurodevelopmental disorders (Durand et al., 2012; Yashiro et al., 2009). In an effort to understand the mechanism underlying this effect, we have discovered that synaptic upscaling is exaggerated in *Ppt1^-/-^* neurons both *in vitro* (**Figure 3**) and *in vivo* (**Figure 6**) and that this contributes to disrupted excitatory synapse formation in the visual cortex of developing mice. In addition to biochemical and immunofluorescence data showing a heightened membrane incorporation of GluA1 in upscaled *Ppt1^-/-^* neurons, calcium imaging data show an increased contribution of NASPM-sensitive, CP-AMPARs to spontaneous activity in dissociated *Ppt1^-/-^* neurons (**Figures 4**). Synaptic upscaling is associated with an increase in GluA1 palmitoylation solely in *Ppt1^-/-^* neurons (**Figure 5**). Palmitome profiling revealed an over-palmitoylation of numerous synaptic proteins in the *Ppt1^-/-^* visual cortex at a young age (∼1.5 months) but pointed to the scaffolding protein Akap5 and associated signaling molecules as collectively showing increases in the palmitoylated fraction (**Figures 7**). Correspondingly, the NFAT transcriptional pathway is sensitized in *Ppt1^-/-^* neurons in response to stimulated neuronal activity and *in vivo* induction of synaptic scaling exacerbated microgliosis (**Figure 8**). Our results indicate loss of lysosomal depalmitoylation disrupts proteostasis of CP-AMPARs and Akap5, thereby exaggerating synaptic upscaling and leading to neuroinflammation.

Interestingly, although we did not pursue it in detail here, our biochemical and immunolabeling assays show that surface expression of GluA1 is not reduced in *Ppt1^-/-^* neurons in response to bicuculline treatment (**Figure 3**), suggesting a failure to scale down synapses. However, further studies are required to determine this effect and is beyond the scope of the current study.

### Dark Rearing Exacerbates CLN1 Pathology: Contribution of Exaggerated Synaptic Upscaling

The current study employed DR as a model of synaptic scaling of the visual system *in vivo* (Desai et al., 2002b; Goel and Lee, 2007a; Goel et al., 2006a, 2011). Intriguingly, DR-*Ppt1^-/-^* mice show exacerbated CLN1 pathology (**Figure 1**). In the case of monocular deprivation in rodents, another method of visual deprivation, both Hebbian and homeostatic plasticity play a role in the short- and long-term visual cortical response (Desai et al., 2002b; Gordon and Stryker, 1996; Keck et al., 2017; Rittenhouse et al., 1999). In the DR paradigm, Goel et al., showed an induction of CP-AMPAR upscaling (Goel and Lee, 2007b; Goel et al., 2006b), and we find that DR associated with this form of synaptic scaling accelerated AFSM accumulation. In contrast, one would anticipate that an LTD-like phenomenon may reduce AFSM load and slow the progression of CLN1 symptoms. For example, in a mouse model of Angelman syndrome that showed low neuronal plasticity at baseline, DR prolonged the lifespan and improved symptoms (Yashiro et al., 2009).

We previously demonstrated that the developmental GluN2 subunit shift is disrupted in in *Ppt1^-/-^* neurons, partly because of over-palmitoylation of GluN2B (Koster et al., 2019). Here, we show this finding persists *in vivo* (**Figure S2C**). N-methyl-D-aspartate receptor (NMDAR) composition significantly affects the calcium dynamics and circuit formation of visual cortical neurons and, importantly, DR is known to cause the persistence of GluN2B at the synapse in visual cortex (Philpot et al., 2001). Therefore, although synaptic upscaling of CP-AMPARs exacerbated the pathology in *Ppt1^-/-^* mice, the bias toward GluN2B-containing NMDARs in *Ppt1^-/-^* visual cortex and their persistence in DR animals may also contribute to the worsened symptoms in DR-*Ppt1^-/-^* mice.

In fact, it is plausible that these mechanisms are intertwined in the *Ppt1^-/-^* brain. It is established that NMDAR activity in the developing brain suppresses unsilencing of synapses (Adesnik et al., 2008) and that GluN2B-NMDAR activity specifically maintains silent synapses (Gray et al., 2011; Hall et al., 2007), indicating the developmental shift from GluN2B- to GluN2A-containing receptors is crucial for the conversion of silent to functionally mature synapses (Hanse et al., 2013). As mentioned, this process is stagnated in the *Ppt1^-/-^* visual cortex and accompanied by an overabundance of thin, filopodial spines in *Ppt1^-/-^* neurons (Koster et al., 2019), which are characteristics of silent synapses (Feldman, 2009) and represent open slots for AMPAR insertion following a strong stimulus, such as induction of synaptic upscaling. Therefore, we postulate that at baseline, where we see a decrease in the frequency of AMPAR-mediated ESPCs in *Ppt1^-/-^* neurons, synapses are immature, dominated by GluN2B-containing NMDARs and remain silent beyond what occurs in typical neurodevelopment. This also suggests a larger population of extrasynaptic AMPARs that are can be mobilized in response to a robust stimulus. Importantly, the increased number of filopodial protrusions in *Ppt1^-/-^* neurons act as sites for scaling-stimulated AMPAR insertion, thereby allowing for an increased capacity of synaptic upscaling (**Figure S4**). Intriguingly, a mouse model of Fragile X syndrome, another neurodevelopmental disorder that presents with intellectual disability and seizure in humans, demonstrates increased dendritic spine density and persistence of silent synapses in the primary somatosensory cortex, echoing our results (Harlow et al., 2010). This model also explains the vulnerability of *Ppt1^-/-^* neurons to excitotoxicity induced by NMDA but not AMPA (Finn et al., 2012).

### Disrupted Depalmitoylation-Dependent Proteostasis Results in Excessive Synaptic Upscaling in *Ppt1^-/-^*Neurons

One of the major findings of the current study is that synaptic upscaling of CP-AMPARs is exaggerated in *Ppt1^-/-^* neurons. There are two primary explanations from our data; 1) over-palmitoylation of GluA1 itself causes its increased surface expression during upscaling or 2) overly palmitoylated Akap5 disrupts the physiological turnover of CP-AMPARs, resulting in their surface retention during scaling.

An initial study describing the phenomenon of synaptic scaling by O’Brien and colleagues showed that the synaptic accumulation of GluA1-containing AMPARs was partly attributable to an increase in half-life of GluA1 protein (O’Brien et al., 1998). Gorenberg et al., recently reported that the extracellular cysteine, C323, is likely to be the site of depalmitoylation by Ppt1 (Gorenberg et al., 2020). The authors speculated that the C323 palmitoylation site represents a latent signal, whereby unmodified GluA1 is palmitoylated at C323 during trafficking to the synapse but forms a disulfide bond with C75 upon exposure to the synaptic cleft. Though it is not yet known how palmitoylation at this site affects GluA1 trafficking, we argue that lack of depalmitoylation by Ppt1 results in the excessive palmitoylation of newly synthesized GluA1 during upscaling and leads to its increased retention at the membrane. It is also established that palmitoylation influences the turnover of specific proteins by inhibiting their ubiquitination (Linder and Deschenes, 2007), either directly or by limiting their endocytosis and exposure to ubiquitination enzymes prior to lysosomal degradation. GluA1 ubiquitination increases during synaptic downscaling (Jewett et al., 2015; Scudder et al., 2014). Consequently, one possibility is that increased GluA1 palmitoylation in upscaled *Ppt1^-/-^* neurons reduces its degradation and promotes a synaptic accumulation of CP-AMPARs.

Alternatively, the aberrant trafficking of CP-AMPARs during synaptic scaling may not arise from palmitoylation of GluA1 directly, as we also observe excessive palmitoylation of Akap5 and its associated signaling molecules in *Ppt1^-/-^* cortex. A series of experiments implicates Akap5 in the regulation of CP-AMPARs during synaptic scaling (Sanderson et al., 2018). These authors show that palmitoylated Akap5 limits the basal synaptic incorporation of CP-AMPARs but is required for their postsynaptic insertion during LTP (Purkey et al., 2018). Further, Akap5 requires depalmitoylation for its removal from the postsynaptic site and undergoes ubiquitination to downregulate GluA1-containing AMPARs during chemical LTD (Cheng et al., 2020; Woolfrey et al., 2018). Therefore, we anticipate that Akap5 normally undergoes depalmitoylation-dependent degradation (Cheng et al., 2020) and that this mechanism is diminished in *Ppt1^-/-^* neurons. Consequently, we postulate that overly-palmitoylated Akap5 harbors an enlarged perisynaptic pool of CP-AMPARs that is mobilized during synaptic upscaling in *Ppt1^-/-^* neurons either at the extrasynapse or dendritic endosomes.

Lastly, we demonstrate that the over-palmitoylation of Akap5 leads to a sensitization of the NFAT transcriptional program in *Ppt1^-/-^* neurons (**Figure 8**). Not only does this likely represent a link to the induction of neuroinflammation that is a predominant feature of CLN1 (Anderson et al., 2013; Radke et al., 2015) and CLN1 mouse models (Jalanko et al., 2005; Kielar et al., 2007; Macauley et al., 2009, 2011, 2014), but a target of NFAT in neural cells is TNFα (Canellada et al., 2006). TNFα is required for synaptic scaling (Stellwagen and Malenka, 2006), acting as a permissive signal to maintain a plastic state that allows for scaling of postsynaptic responses (Steinmetz and Turrigiano, 2010). Furthermore, the calcineurin-NFAT pathway mediates synaptic scaling through the turnover of CP-AMPARs (Kim and Ziff, 2014). Therefore, it is plausible that mis-localization of Akap5 or impaired interactions with calcineurin drive over-activation of NFAT and transcription of TNFα, permitting an exaggerated scaling response of CP-AMPARs in the *Ppt1^-/-^* brain (**Figure S4**). Therefore, several related mechanisms may collaborate to drive exaggeration of CP-AMPAR incorporation during synaptic scaling up in *Ppt1^-/-^* neurons.

### Implications for CLN1 Progression and Therapeutic Intervention: Two Novel Therapeutic Strategies

The function of Ppt1 in neurons remains incompletely characterized and understanding its role at the synapse is of critical importance to identifying and eventually correcting pathophysiology in CLN1. Typically, sensory cortical networks desynchronize during development, likely expanding the information encoding capability of the neuronal network (Golshani et al., 2009; He et al., 2018; Rochefort et al., 2009). In our previous study, we show that a developmental maturation of NMDA receptors is disrupted in the visual cortex, favoring the synaptic incorporation of GluN2B, which is associated with an increase in the neuronal plasticity threshold (Abraham, 2008; Yashiro and Philpot, 2008). The current study adds another layer of synaptic dysregulation and shows that synaptic upscaling of CP-AMPARs is exaggerated in *Ppt1^-/-^* visual cortical neurons. Interestingly, in a computational study, artificial deafferentiation of a population of synthetic neurons drives a pathological synchrony in their activity due to synaptic scaling (Fröhlich et al., 2008). Therefore, the disruption of these molecular synaptic plasticity mechanisms likely contributes to the network hypersynchrony we observe in the *Ppt1^-/-^* visual cortex (**Figure 6**). Whether this manifests in *Ppt1^-/-^* visual cortex as a failure to desynchronize during development, or a resynchronization because of disrupted plasticity remains unclear. Nevertheless, we postulate that this pathological network activity eventually contributes to seizure activity in *Ppt1^-/-^* mice. Indeed, this failure to desynchronize is also observed in Fragile X syndrome (Gonçalves et al., 2013). Taken together, we propose a model by which disrupted NMDAR and AMPAR proteostasis underlies network desynchronization, driving excitotoxic cell death and synaptic scaling of the afferent brain region. A similar framework has been proposed to contribute to the pathogenesis of Alzheimer disease (Small, 2008). In this way, sequential scaling- synchronization cycles leads to a spreading neurodegeneration in parallel with seizure activity, as observed in CLN1.

Intervention at the molecular level to improve this synchronous oscillatory activity, which has been recently performed by targeting NMDA receptors in models of Alzheimer disease (Hanson et al., 2020), may prove beneficial in CLN1. On the other hand, if the consequence of Ppt1 mutation is a loss of developmental desynchronization, perhaps a therapeutic strategy targeted at the circuit level, such as deep brain stimulation or emerging technologies that can correct network activity (Rowan et al., 2014; Tye, 2014) might prove beneficial. At minimum, our findings related to the dysregulation of CP-AMPARs (GluA1) in particular lend credence to the recent clinical use of Perampanel as a second line treatment for CLN1 (Augustine et al., 2021).

In parallel to the dysregulated homeostatic plasticity, our data suggest that the over-palmitoylation of Akap5 contributes to the pathogenesis of the *Ppt1^-/-^* brain through its regulation of CP-AMPAR proteostasisand downstream signaling through the calcineurin-NFAT axis. This finding links aberrant calcium activity to a neuroinflammatory cascade and gliosis in CLN1 for the first time (**Figure 8**). Further, Calcineurin-NFAT activity represents a new therapeutic target in CLN1 that has a history of being inhibited in other conditions requiring immunosuppression with FDA-approved drugs such as Tacrolimus (FK506) and cyclosporin. Interestingly, Tacrolimus is currently being tested in a phase 2 clinical trial for treatment of Alzheimer disease (trial: NCT04263519), suggesting this or next generation analogs may be a promising intervention in multiple neurodegenerative diseases. Future studies focused on testing the efficacy of these (or other) calcineurin- directed therapeutics will be of great interest to the CLN1 field and patient population.

In conclusion, we demonstrate here with an interdisciplinary approach how failed protein depalmitoylation disrupts synaptic plasticity via impaired proteostasis and triggers neuroinflammatory signaling that underlies a devastating disease and offer multiple novel targets for therapeutic intervention.

## Author contributions

Conceptualization, A.Y., K.Y.T., S.M.C., and K.P.K; Methodology, K.P.K; Formal analysis, K.P.K., T.A.N.; Investigation, K.P.K, E.F.B., E.A.V., T.T.A.N., A.N., L.N., Z.F.; Resources, K.P.K.; Writing – Original draft, ., A.Y.; Writing – Review and editing, K.P.K., A.Y., K.Y.T., S.M.C.; Visualization, K.P.K.; Supervision, A.Y., K.Y.T., S.M.C.; Funding acquisition, A.Y.

## Conflict of interest statement

The authors declare no conflicts of interest.

## Experimental Procedures

### Animals, group allocation, and data handling

All animal procedures were performed in accordance with the guidelines of the University of Illinois of Chicago Institutional Animal Care and Use Committee. *Ppt1^+/-^* (heterozygous) mice were obtained from Jackson Laboratory and maintained on 12h light/dark cycle with food and water *ad libitum.* Breeding of *Ppt1^+/-^* animals results in litters containing *Ppt1^-/-^*, *Ppt1^+/-^*, and *Ppt1^+/+^* (WT) animals. *Ppt1^-/-^* and WT littermate controls were genotyped in-house (Gupta et al., 2001) and used for experiments at specified developmental time points: P11, P14, P28, P33, P42, P60, P78, and P120. Though we used the littermate control system, in which WT and *Ppt1^-/-^* mice from the same litters were compared, each n was treated independently in statistical testing for *in vivo* comparisons (pair-wise tests were not used). In contrast, *in vitro* and imaging data were treated as paired analyses, since each culture was often collected, processed, or immunostained at distinct times (rather than multiple n being processed simultaneously). Imaging data was acquired randomly for each experiment (no criteria for selecting cells, view fields, etc. except where anatomically necessary). All data was acquired and maintained without descriptive naming/labeling to ease randomization. Data was either analyzed by lab members blinded to condition or randomized prior to analysis by KPK, with the exception that 2-photon data was not randomized before analysis; however, these calcium data are extracted in a fully automated manner by the EZ calcium Matlab plugin (see below).

### Brain fractionation, biochemical assays from tissue samples, and immunoblotting

For collection of brain for biochemistry (immunoblot), *Ppt1^-/-^* and WT animals were decapitated following isoflurane anesthesia, then the brain was removed, and washed in ice-cold PBS. The occipital cortex (visual cortex), hippocampus, and remaining cortex were dissected and separately collected on an ice block. Isolated visual cortices from *Ppt1^-/-^* and WT animals were homogenized in ice-cold synaptosome buffer (320mM sucrose, 1mM EDTA, 4mM HEPES, pH7.4 containing 1x protease inhibitor cocktail (Roche), 1x phosphatase inhibitor cocktail (Roche) and 1mM PMSF) using 30 strokes in a Dounce homogenizer. Aliquots of lysates were stored at -80°C and the remaining sample was used for synaptosome preparation, performed as previously with slight modification. In brief, WLs were centrifuged at 1,000 x g to remove cellular debris, supernatant was then centrifuged at 12,000 x g for 15min to generate pellet P2. P2 fraction was resuspended in synaptosome buffer and spun at 18,000 x g for 15min to produce synaptosomal membrane fraction, LP1, which was used for downstream biochemical analyses (synaptosomes). For immunoblot, protein concentration of each sample was determined using BCA protein assay (Pierce). Samples were then measured to 20μg total protein in 2x Laemmli buffer containing 10% b-mercaptoethanol (Bio-rad), heated at 70°C for 10min and loaded into 4-20% precast gels (Bio-rad) for electrophoresis (130V, 1.5-2h). Proteins were wet-transferred to PVDF membranes (Immobilon-P, Millipore), blocked in TBS, pH7.4 containing 5% non-fat milk and 0.1% Tween-20 (TBS-T+5% milk). Membranes were incubated in primary antibody solutions containing 2% BSA in TBS-T for 2h at room temperature (RT) or overnight at 4°C. Primary antibodies were used according to the **Key Resources Table**. Membranes were then incubated with appropriate secondary, HRP-conjugated antibodies (Jackson ImmunoResearch) at either 1: 1,000 or 1:5,000 for 1h at RT before washing three times with TBS-T. Visualization and quantification was performed using Pierce SuperSignal ECL substrate and Odyssey-FC chemiluminescent imaging station (LI-COR). Signal density for each synaptic protein was measured using the LI-COR software, Image Studio Lite (version 5.2) and was normalized to the signal density for β-actin loading control for each lane. The number of visual cortices used for analysis at each age are displayed in Figure 1 (individual points on graph), with two technical replicates for each experiment averaged together.

### APEGS assay from visual cortices

The APEGS assay was performed as described by Kanadome and colleagues (Kanadome et al., 2019), following the guidelines for tissue samples. Visual cortices first underwent the synaptosome preparation protocol as above, except that homogenate buffer used was as directed by the APEGS protocol (20mM Tris-HCl, 2mM EDTA, 0.32M sucrose, pH 8.0). Lysates and synaptosomes were then brought to 300μg total protein in a final volume of 0.5ml buffer B (PBS containing 4% SDS, 5mM EDTA, 8.9M urea, and protease inhibitors). The remaining sample was used for inputs. 300μg protein was reduced by addition of 25mM Bond-Breaker™ TCEP (0.5M stock solution, ThermoFisher) and incubation at RT for 1h. Next, to block free thiols, freshly prepared N-ethylmaleimide (NEM) in 100% ethanol was added to lysates (to 50mM) and the mixture was rotated end-over-end for 3h at RT. Following 2x chloroform-methanol precipitation (at which point, protein precipitates were often stored overnight at -20°C), lysates were divided into +hydroxylamine (HA) and –HA groups for each sample, which were exposed to 3 volumes of HA-containing buffer (1M HA, to expose palmitoylated cysteine residues) or Tris-buffer control (-HA), respectively, for 1h at 37°C. Following chloroform-methanol precipitation, the samples were solubilized and exposed to 10mM TCEP and 20mM mPEG-5k (Laysan Bio Inc, see **Key Resources Table**) for 1h at RT with shaking (thereby replacing palmitic acid with mPEG-5K on exposed cysteine residues). Following the final chloroform-methanol precipitation, samples were solubilized in a small volume (70μl) of PBS containing 1% SDS and protein concentration was measured by BCA assay (Pierce). Samples were then brought to 20μg protein in Laemmli buffer with 2% β-mercaptoethanol for immunoblot analyses as above. Quantification of palmitoylated vs. non-palmitoylated protein was carried out as for standard immunoblot analysis, with the additional consideration that signal from palmitoylated bands demonstrating the APEGS-dependent molecular weight shift was divided by the signal from the non-palmitoylated band, the location of which was verified by matching to the –HA control sample. This ratio was divided by β-actin control from the same lane for normalization.

### Acyl-biotin exchange (ABE) assay for palmitoyl-proteomics

WT and *Ppt1^-/-^* occipital cortices from P42 were used for palmitoyl-proteomic analysis. Lysates and synaptosomes were collected as described above. The palmitoyl-proteomic protocol was then carried out according to Wan et al., (2007) with slight modifications (Wan et al., 2007). Before beginning the assay, a BCA assay was performed to start with equal (600μg) protein content for each sample. Blocking (N-ethylmaleimide; NEM), hydroxylamine, and biotinylation (HPDP-biotin) steps were all performed as recommended in the protocol. The elution protocol was also followed (Wan et al., 2007), with the exception that instead of streptavidin resin, magnetic streptavidin coated beads (Dynabeads™, ThermoFisher) were used (100μl beads/sample). The final eluent was frozen at -80⁰C and prepared for mass spectrometry (see below). Due to the small starting material (occipital cortex only), the whole procedure was scaled down into 2mL tubes, including chloroform-methanol precipitations, which were carried out with the following volumes: 150μl sample, 600μl methanol, 150μl chloroform, 450μl water. This limited protein loss, which was evident in trial runs using 15ml conical tubes.

### Mass spectrometry

Eluents from the ABE assay were dried for approximately 30 minutes and digested using S-trap Micro Spin Column Digestion protocol (Protifi, Huntington, NY) with minor changes. Briefly, 30 µL of 10% sodium dodecyl sulfate (SDS) 100 mM triethylammonium bicarbonate (TEAB) with Pierce protease inhibitor cocktail (ThermoFisher Scientific, Waltham, MA) and phosphatase inhibitors (10 mM sodium pyrophosphate, 1 mM PMSF, 1 mM sodium orthovanidate, and 1 mM β-glycerolphosphate). Proteins were reduced with 20 mM final concentration of dithiothreitol (DDT) at 95 °C for 10 minutes, followed by alkylation in the dark, at room temperature, with 40 mM of iodoacetamide. Next, phosphoric acid was added for a final concentration of 1.2%. Samples were briefly vortexed to mix before 300 µL of S-trap binding buffer (90% MeOH, 100 mM TEAB) was added. Samples were vortexed again prior to loading onto the S-Trap Micro Spin Columns. After four washes with 150 µL of S-trap binding buffer with centrifugation at 1,000x g, 40 µL of 50 mM TEAB containing 0.75 µg of trypsin was added and incubated overnight at 37 °C.

Peptides were eluted with 40 µL of each of the following solutions: 50 mM TEAB, 0.2 % formic acid (FA), and 50% acetonitrile (ACN) 0.1% FA. Spin column was spun at 4,000x g in after adding each solution. Pooled eluents were dried down prior to resuspension in 100 µL of 3% ACN 0.1% FA.

### LC-MS Analysis

Three µL of resuspended samples were injected for LC-MS/MS analysis, similar to a previously mentioned method (Nguyen et al., 2019). Briefly, peptides were loaded onto a Thermo NanoViper trap column (75 µm x 20 mm, 3 µm C18, 100 Å) (Thermo Fisher Scientific, Bremen, Germany) using an Agilent 1260 Infinity nanoLC system (Agilent Technologies, Santa Clara, CA) and washed for 10 minutes with 0.1% FA at 2 µL/min. Peptides were separated with a 120-minute gradient (from 5-60% ACN with 0.1% FA), at 0.25 µL/min flow rate, on an Agilent Zorbax 300SB-C18 column (75 µm x 150 mm, 3.5 µm, 300 Å). Data was collected using data-dependent acquisition (DDA) analysis by a Thermo Q Exactive mass spectrometer (Thermo Fisher Scientific, Bremen, Germany). Settings for the mass spectrometer are as follows: capillary temperature at 250 °C, spray voltage 1.5 kV, MS1 scan at 70,000 resolution, scanning from 375-1600 m/z, automatic gain control (AGC) target 1E6 for a maximum injection time (IT) of 100 ms. The ten most abundant peaks within an MS1 spectrum were isolated for MS/MS, with an isolation width at 1.5 m/z and dynamic active exclusion set for 20 s. MS/MS spectra were collected at 17,500 resolution, for a maximum of 50 ms or a minimum of 1E5 ions. Normalized collision energy (NCE) was set at 27%. Masses with charges of 1 and larger than 6 were excluded from MS/MS analysis. **Analysis for palmitoyl-proteomics**

Raw files were searched with Proteome Discoverer 2.3 (Thermo Fisher Scientific, Waltham, MA) using the Sequest HT search engine against the UniProt *Mus musculus* database (22,286 gene sequence; downloaded April 27, 2017). Mass error tolerance was set to 10 ppm for precursors, cleaved by trypsin, allowing a maximum of two missed cleavages, with sequence lengths between 6 and 144 amino acids. Fragment masses were searched with a tolerance of ± 0.02 Da. Dynamic modifications included oxidation (M), deamidation (N, R, Q), and acetylation (N-terminus). Carbamidomethylation was set as a static modification (C). Both peptides and PSMs were set to a target false discovery rate (FDR) ≤ 0.01 for matches with high confidence. Label-free quantification (LFQ) was performed using precursor ion intensity. Samples were normalized using the average intensity of all peptides. The top 5 most abundant peptides were used for protein abundance calculation. t-test were used to determine the p-values between the two conditions.

Datasets filtered for 2 unique peptides were further narrowed by filtering for proteins showing an increase in their abundance ratio (*Ppt1^-/-^/*WT) of >1.2 fold. These filtered gene lists for lysates and synaptosomes were input separately into the SynGO online tool (Koopmans et al., 2019). The genes encompassed in the top significant biological process (BP) SynGO-term for lysates and synaptosomes were then input into STRING, the online protein-protein interaction database (https://string-db.org/). A K-means cluster analysis was performed to detect clusters of functionally related proteins and for clarity of visualization.

### Transcardial perfusion, immunohistochemistry, and image analysis

*Ppt1^-/-^* and WT mice were anesthetized using isoflurane and transcardially perfused with ice cold PBS (pH 7.4, ∼30ml/mouse) followed by 4% paraformaldehyde (PFA) in PBS (∼15ml/mouse). Brains were removed and post-fixed overnight at 4°C in 4% PFA and transferred to PBS, pH7.4 containing 0.01% sodium azide for storage if necessary. Brains from *Ppt1^-/-^* and WT animals were incubated in 30% sucrose solution for 48h prior to sectioning at either 50 or 100μm in cold PBS using a Vibratome 1000 (Technical Products International, St. Louis, MO). Serial sections were stored free floating in cryoprotectant solution (30% glycerol, 30% ethylene glycol in PBS) at -20°C until analysis of AFSM or immunohistochemistry was performed.

For AFSM analysis (as in (Koster et al., 2019), 3-4 mid-sagittal sections were mounted on Superfrost Plus microscope slides (VWR) using Vectamount mounting media containing DAPI (Vector Laboratories). Images were acquired for at least 2 sections from each animal using a Zeiss LSM710 confocal laser scanning microscope at 40x magnification (excitation at 405nm to visualize DAPI and 561nm to visualize AL). All sections were imaged using identical capture conditions. Quantification of AFSM was performed by thresholding images in FIJI (NIH), generating a binary mask of AFSM-positive pixels (satisfied threshold) vs. background. The identical threshold was applied to each image (from cortical surface to subcortical white matter and across animals). Percent area occupied by AFSM puncta that satisfied the threshold was then calculated using the “analyze particles” tool in FIJI. This analysis was performed for 2-4 sections (total of ∼10-20 images, as imaging an entire cortical column is typically 5 interlaced images) from each animal and averaged together to give a single value, representative of the total area occupied by AFSM in the cortical column imaged. Three to six animals per group were analyzed this way and averaged to give the mean area occupied by AFSM at each time point, for both genotypes (n=3-6 animals/group).

For immunohistochemistry, 3-4 slices medial sections were first incubated in TBS for 10 min before undergoing permeabilization (TBS + 0.5% Triton X-100) for 30 minutes at RT. Next, samples underwent antigen retrieval by heating in tris-EDTA (pH 9.0) at 95°C for 30 minutes before being equilibrated to room temperature in tris-EDTA solution for 40 minutes. Tissue was then blocked (TBS + 0.1% Triton X-100, 4% BSA, and 5% normal goat serum) for 2 hours at RT before being incubated in rabbit anti-Iba1 (1:1,000, in TBS + 0.1% Triton X-100 + 2% BSA) for 48 hours. After washing 4 times, 10 minutes each, tissue was incubated in Alexa Fluor 488 goat anti-rabbit (1:1,000; ThermoFisher) overnight at RT. The tissue was then washed (4x, 10 minutes) and mounted as above.

Microglial images were acquired with a Zeiss LSM710 confocal microscope either at 10x (low magnification images) or 63x for Sholl analysis. 63x images were acquired as 30-60 plane Z-stacks (Z-interval = 2μm) in random fashion (the only criterion is that at least one full microglia had to be centered in the stack) at the border of layer 2/3 and 4 in the visual cortex. 3-4 images were taken from 2 tissue sections for each animal. Images were analyzed using the Sholl analysis tool in Fiji by lab members blinded to the condition. Briefly, the image was collapsed into a maximum-intensity projection to ensure all microglia processes were captured and individual microglia were outlined with a freehand ROI. The surrounding area was removed (“clear outside” function in Fiji) and the image was then thresholded to generate a mask of all microglia processes and an ROI was created at the center of the cell soma. The mask containing 1 individual microglia was then skeletonized using the “skeletonize” plugin in Fiji. Sholl analysis was performed according to these parameters using the Fiji Sholl tool: start radius (from the center of cell soma ROI) = 5μm, step size = 2μm, end radius = 70μm. The number of intersections at each 2μm step was averaged for all microglia from each animal and counted as one n.

## Electrophysiology

WT and *Ppt1^-/-^* animals at P15 or P28-32 were deeply anesthetized via intraperitoneal injection of chloral hydrate and decapitated. All recordings were conducted from layer 2/3 pyramidal neurons of monocular visual cortex (350-μm-thick coronal slices) at 33-35°C using a low-chloride-based internal solution and an external solution free of glutamate and GABA blockers to enable concurrent acquisition of excitatory and inhibitory synaptic currents at the single-cell level. Contents of the internal solution are the following (in mM): 10 CsCl, 130 Gluconic acid, 10 HEPES, 2 MgCl_2_, 5 NaATP, 0.6 NaGTP, and 3 QX-314, pH 7.23-7.28, 280-282 mOsm. The recording artificial cerebral spinal fluid contained the following (in mM): 122.5 NaCl, 3.5 KCl, 25 NaHCO_3_, 1 NaH_2_PO_4_, 2.5 CaCl_2_, 1 MgCl_2_, 20 glucose, and 1 ascorbic acid, pH 7.40–7.43, 295–305 mOsm. As a result, both spontaneous glutamatergic and GABAergic events could be assessed readily by recording the frequency of postsynaptic currents (PSCs) at the −60 mV (PSC_−60 mV_) and +15 mV (PSC_+15 mV_) holding potentials, respectively. Only neurons with at least 10 min of stable baseline activity were included for analyses. The frequency of PSC_−60mV_ and PSC_+15mV_ events from at least two noncontiguous epochs of 60 s each was compared.

### Stereotaxic viral injection

Injection into neonatal mouse cortices was performed according to He et al., (2018). P0 or P1 pups underwent hypothermia-induced anesthesia (incubation on an ice block for 5-10min) before being placed in the mouse stereotaxic frame (on a second ice block) equipped with specialized ear bars for holding pups (RWD Life Science Inc). AAV.CamKII.GCaMP6f.WPRE.SV40 (AAV9) (Addgene) and pAAV-hSyn-mCherry (AAV2) were pre-mixed in 1:1 ratio before being backloaded into a Neuros syringe (33 gauge, Hamilton). Alternatively, a Nanoject 3 automatic injector (Drummond Scientific Company; Broomall, PA) was used with a finely pulled borosilicate capillary micropipette. 200nL of the mixture was injected at a 15° angle approximately 150μm below the dura surface to transfect a large population of layer 2/3 excitatory cortical neurons with the viral particles. The needle was left in place for 1 minute, then lifted by a minimal distance and left in place for an additional 30 seconds before being slowly removed to avoid reflux of solution along the needle track. Pups were immediately placed on a heating pad and monitored until they regained consciousness.

For DREADD-induced scaling experiments, animals between P16-18 were anesthetized via isoflurane inhalation (4% induction, 1-1.5% maintenance) and placed in a stereotaxic frame (RWD Life Science Inc.). A small (0.5-1cm) incision was made to expose the posterior portion of the skull. Connective tissue beneath the scalp was removed using sterile cotton swabs and the coordinates of bregma were recorded. The injection needle was then moved to the appropriate injection location (2.2mm posterior to bregma or ∼0.4mm anterior to lambda and 2.2mm lateral) for the left visual cortex and the location was marked on the skull using a fine tip permanent pen. A small burr hole was drilled in this location with a dental drill (Marathon) and the needle was lowered so that it rested just above the cortical surface. The needle was then lowered into the cortex (400μm depth) and 50nl of virus was ejected. The needle was then slowly retracted at 100μm increments, and 50nl of virus was injected at each increment (200nl total through dorsoventral aspect of the visual cortex). The same procedure was performed for animals in which the control (hDlx-GFP) virus was injected. After removal of the needle (10min total), the scalp was sutured and animals were monitored until recovery, at which point the were returned to their home cages.

Following a recovery period of ∼10 days to allows for expression of the DREADD in visual cortical neurons, animals were injected with the DREADD agonist, CNO (3mg/kg) daily for five consecutive days, and sacrificed for biochemical analysis 4-6 hours after the final injection.

### Cranial window implantation

Cranial window implantation was performed as previously described with some modification (Holtmaat et al., 2009). Virus injected animals were anesthetized via isoflurane inhalation (4% induction, 1-1.5% maintenance) and placed in a stereotaxic frame (RWD Life Science Inc.). All procedures were performed under sterile conditions. Buprenorphine-SR (1mg/kg, subcutaneous), meloxicam (2mg/kg, subcutaneous), and dexamethasone (2mg/kg, subcutaneous) were administered preoperatively. Hair from the scalp was removed using commercially available Nair by applying it with a cotton swab, waiting 3 minutes, and removing the hair with several clean cotton swabs. The scalp was then cleaned by sequential application of betadine-iodine (3x, allowing to dry in between each fresh application) and alcohol using alcohol pads (2x). Once the scalp was clean, an approximately 2cm incision was made along the posterior aspect of the midline and the margins of the incision were expanded to visualize the skull covering the left visual cortex by gently pushing aside the scalp tissue with sterile, fine tipped cotton swabs. Once the incision was large enough and centered on the left visual cortex, the connective tissue beneath the scalp were removed from the skull by applying of ∼25μl epinephrine-lidocaine (via insulin syringe) and rubbing with sterile cotton swabs. The margins of the incision were also simultaneously dried with the swab. Once the margins of the tissue were dried, a layer of cyanoacrylate was applied to the edges and cured using minimal application of Zip Kicker cyanoacrylate catalyst (Zap). Next, the position of the headplate (CP-1, Narishige) was marked by positioning it so that the center was atop the visual cortex and marking its location with a fine tip marker. A layer of cyanoacrylate was then applied over the entire surface of exposed skull, leaving sufficient area (∼6mm) so that glue does not cover the eventual drilling site, and the headplate was quickly secured with the glue to the desired position. After at least 5 minutes, allowing for the glue to cure, a 4- 5mm circular piece of skull was removed by drilling around the circle circumference using a dental drill (Marathon) and frequent soaking with sterile PBS to soften the skull. Once the drill site was thin enough, fine forceps were used to lift the medial edge and the circular bone fragment with a diameter of 4mm was removed completely. The exposed brain tissue was covered with PBS-soaked sterile gelfoam (Surgifoam) to clear away any blood, which was minimal. Lastly, a 5mm coverslip was glued to the top of the burr hole on the headplate. Animals were then immediately moved to a heating pad and monitored until they regained consciousness. Animals typically recovered fully within 15 minutes and showed little impact of the headplate implantation on normal functions such as eating and drinking (without any weight loss).

### Cholera toxin B subunit injection

Mice between P20-P40 were anesthetized via isoflurane inhalation (4% induction, 1-1.5% maintenance) and placed in a stereotaxic frame (RWD Life Science Inc.). After expressing a small amount of vitreous from the eye, intravitreous injections of 2μl fluorophore conjugated CTB (1mg/ml; 488nm conjugate in left eye, 555nm conjugate in right eye) were performed for each mouse. Following the injection, antibiotic ointment was applied to each eye and animals were allowed to recover in their home cage for 24h before undergoing transcardial perfusion as detailed above. Sectioned brains (100μm sequential coronal sections encompassing the entire dLGN) were then assessed for the quality of the injections by a blinded researcher and only those brains with well-traced retinogeniculate projections were immunostained for Iba1. Matched sections were then imaged under high magnification at the border of converging retinal projections from both eyes in each dLGN using an LSM710 confocal microscope and compared for microglial morphology (Sholl analysis) as described above.

### Primary cortical neuron culture

For primary cortical neuron cultures, embryos from timed-pregnant, *Ppt1^-/+^* dams were removed, decapitated under anesthesia, and cortices resected at embryonic day (E) 15.5. All dissection steps were performed in ice cold HBSS, pH7.4. Following cortical resection, tissue from each embryo was individually collected to a separate microtube, genotyped, and digested in HBSS containing 20U/ml papain and DNAse at 37°C (20min total, tubes flicked at 10min) before sequential trituration with 1ml (∼15 strokes) and 200μl (∼10 strokes) pipettes, generating a single-cell suspension. For live-cell imaging experiments, cells were counted then plated at 150,000-180,000 cells/well in 24-well plates containing poly-D-lysine/laminin-coated coverslips. For biochemical experiments, i.e. immunoblot, APEGS assay *in vitro,* cells were plated on poly-D-lysine/laminin-coated 6-well plates at 1,000,000 cells/well. Cells were plated and incubated at 37°C in plating medium (Neurobasal medium containing B27 supplement, L-glutamine and glutamate) for 3-5 DIV, before replacing half medium every 3 days with feeding medium (plating medium without glutamate). For synaptic scaling experiments, neurons were treated with either bicuclline (20mM, solubilized in DMSO, Tocris) or TTX-citrate (1mM, solubilized in sterile water, Tocris) for 48 hours where indicated.

### Primary cortical neuron harvesting and immunoblotting

Primary cortical neurons from E15.5 WT and *Ppt1^-/-^* embryos were cultured for 7, 10, or 18 DIV prior to harvest for immunoblot or APEGS assay (only DIV18 used for APEGS). To harvest protein extracts for APEGS, cells were placed on ice, washed 2x with ice-cold PBS before addition of 400μl per well of PBS with 4% SDS and protease inhibitor cocktail (as in Yokoi et al., 2016). Cells were incubated and swirled with lysis buffer for 5 minutes, scraped from the plate, triturated briefly, and collected in 1.5ml tubes. Lysates were centrifuged at 20,000g for 15min to remove debris, and the supernatant was collected for biochemical analysis. Immunoblotting analyses were performed as above. APEGS assay *in vitro* was carried out as described in the following section. **APEGS assay on primary cortical neuron lysates**

The APEGS assay was performed as described by Kanadome et al., (2019) and in Koster et al., (2019). The protocol is nearly identical to the description above for *in vivo* samples, with slight modifications as recommended in the protocol by Kanadome et al. (2019). Specifically, the cortical neuron lysates were brought to 300μg total protein in a final volume of 0.5ml buffer A (PBS containing 4% SDS, 5mM EDTA, protease inhibitors; instead of buffer B). Further, the initial protein reduction by TCEP incubation was performed at 55°C for 1h (rather than at RT). Following the final chloroform-methanol precipitation, samples were solubilized in 60μl of PBS containing 1% SDS and protein concentration was measured by BCA assay (Pierce). Samples were then brought to 10μg protein in Laemmli buffer with 2% β-mercaptoethanol for immunoblot analyses as above. Quantification of palmitoylated versus non-palmitoylated protein was carried out as described for *in vivo* samples.

### Biotinylation

Surface biotinlyation assay was performed as in (Ehlers, 2000) and (Yoshii et al., 2013) with some modifications. Briefly, neuron cultures (6-well plates) were cooled on ice to stop membrane trafficking in cold DPBS plus 1 mM MgCl_2_ and 2.5 mM CaCl_2_, then treated with 1.5 mg/ml sulfo-NHS-biotin in DPBS for 30 min to label surface receptors. Unbound biotin was quenched with cold DPBS plus 1 mM MgCl_2_, 2.5 mM CaCl_2_, and 50 mM glycine (rinse, 2 × 3 min). Neurons were then lysed for 5 min in SDS-RIPA buffer (Cell Signaling) with protease inhibitor cocktail (Roche), scraped and the lysed solution was centrifuged at 16,000× g for 15 min at 4°C. Protein concentration of the supernatant was determined and 400 μg of protein was adjusted to a volume of 1ml with 0.1% SDS-RIPA buffer. Biotinylated GluA1 receptor subunits were precipitated with 75μl streptavidin coated magnetic bead (Thermo Scientific) slurry either for 3 hours at RT or overnight at 4°C with end over end rotation. Beads were then washed with the SDS-RIPA buffer twice before brief centrifugation. Pulled down subunits were eluted into Laemmli sample buffer with 2% b-mercaptoethanol by boiling for 6 min and the magnetic beads were removed from solution. Samples were then loaded onto 4-20% Bio-Rad gradient gels as above for immunoblot analysis.

### Transfection, calcium imaging, FRAP, and GFP-NFAT nuclear translocation analyses

To image calcium signals in WT and *Ppt1^-/-^* neurons, cells were transfected at DIV10 with GCaMP3 (see **Acknowledgements**) using Lipofectamine® 2000 (ThermoFisher) according to manufacturer protocol with a slight modification. GCaMP3 DNA construct (∼2μg/μl, added at ∼1μg/well) was mixed with Lipofectamine-containing Neurobasal medium, incubated for 30min to complex DNA-Lipofectamine, equilibrated to 37°C, and added to the cells 250μl/well for 1h. Following incubation, complete medium was returned to the cells.

Cells were grown to DIV16-18 and a subset of cells were treated with TTX (1mM) for 48 hours. Neurons were imaged with constant perfusion (∼2.5ml/min) of Tyrode’s solution without magnesium (imaging medium: 139mM NaCl, 3mM KCl, 17mM NaHCO_3_, 12mM glucose, and 3mM CaCl_2_) using a LSM710 confocal microscope equipped with heated stage. 16-bit videos were acquired at a resolution of 512 x 512 at ∼5 frames per second at 10x magnification (with 3x digital zoom). Baseline acquisition, NASPM perfusion (1 or 10μm), and washout periods were 4 min, 4, min, and 8 minutes, respectively; this corresponds to 1000 video frames, 1000 video frames, and 2000 video frames, respectively. For analysis, since NASPM persists in the bath for some time after infusion, the final 1000 frames (3000-4000) were always used for the washout period (i.e., frames 2000-3000 were excluded from analysis as a transition period between NASPM and washout, since washing is not complete during that time). All active synapses were encompassed with a minimal-sized circular (spines) or rectangular (shaft) ROI manually for each video by lab members blinded to condition and genotype. A typical video resulted in 120-200 ROIs (synapses). The “multi measure” tool was applied to each ROI to generate a ΔF/F_0_ trace for each synapse. The raw fluorescence data for each synapse was then normalized to its own baseline and split into 1000 frame portions corresponding to baseline, NASPM, and washout conditions. The number of active frames for each period were counted by summing the frames reaching a threshold plus 3 x standard deviation (SD) for each synapse. A synapse was considered to be inhibited by NASPM if it met the following criteria: 1) The synapse had to be active, i.e. show at least 6 active frames (+3 x SD) during the baseline period, 2) the synapse showed a reduction in active frames in the NASPM period by more than 80%, and 3) the synapse recovered by showing at least 5 active frames in the washout period. The total activity plots (counted active frames) are plotted in Figure 4C, while the proportion of inhibited synapses (synapses which met the above criteria) is plotted in Figure 4D.

For FRAP analyses, WT and *Ppt1^-/-^* cortical neurons were transfected with SEP-GluA1 (Kopec et al., 2006) at DIV10 as described above. Neurons at DIV16 were imaged in Tyrode’s solution (in mM: 135 NaCl, 5 KCl, 2 CaCl2, 1 MgCl2, 25 HEPES, and 10 glucose, pH 7.4) using a Zeiss LSM880 in Airyscan mode. Five to six spines were randomly selected for each neuron to bleach during the imaging program. Images were acquired every 15 seconds for 5 frames to establish baseline fluorescence before bleaching at maximum laser intensity. Images were then acquired every 15 seconds thereafter for 10 minutes to analyze fluorescence recovery at bleached spines. The Zen Definite Focus function was utilized to correct potential drifting of the imaging plane. To analyze FRAP of SEP-GluA1, manual ROIs were drawn over the bleached spines and one to two control (unbleached) spines for each cell and the ΔF/F_0_ was extracted using the multi measure tool. Baseline fluorescence (average of first 5 frames) was set to 100%, and the following frames were compared to that baseline, generating the FRAP curves in Figure 6. FRAP curves were fit using a non-linear regression in Graphpad Prism 9.0 using the LOWESS function. The immobile fraction was calculated by subtracting the average of the fluorescence recovery value for the final 5 frames for each spine from 100%. All spines for each cell were averaged to create one n. cells were taken from two independent cultures.

Analysis of NFAT nuclear translocation following culture-wide depolarization was performed as in Murphy et al., (2014) with minor modification. Neurons were transfected at DIV 12 with roughly 1:1 mixture of CAG-mCherry and mouse GFP-NFATc3 (pCMV-SGFP2-mNFATc3.dna, courteously provided by Dr. Dell’Acqua), 1mg DNA per coverslip using Lipofectamine® 2000 (ThermoFisher) as above. A subset of neurons was treated with TTX (1μM) for 48 hours leading up to the assay to induce synaptic scaling. The assay was then performed as schematized in Figure 8A. Solutions were composed as follows: Tyrode + TTX (in mM: 135 NaCl, 5 KCl, 2 CaCl2, 1 MgCl2, 25 HEPES, and 10 glucose, pH 7.4 plus 1mM TTX), Depolarization solution was isotonic, but with 50mM KCl (85mM NaCl) and without TTX, and recovery solution was standard Tyrode without TTX. Cells were incubated at 37⁰C in between steps. For every coverslip of depolarized (KCl) neurons, a control with no depolarization (sham, Tyrode solution + TTX) and depolarization with NASPM (KCl + 10 μM NASPM) condition were performed in parallel. Following fixation at the end of the assay, cells were immunostained with anti-GFP 488 antibody (ThermoFisher) following the protocol above to amplify the GFP-NFAT signal. GFP-NFAT nucleus/soma ratio was analyzed by manually tracing the nucleus, based on the DAPI staining signal, and soma as independent ROIs and dividing the integrated fluorescence value (measured by multi measure tool in Fiji) for nucleus by that for the soma.

### Immunocytochemistry

All experimental and control groups were immunostained simultaneously to match exact staining conditions. Neurons were first immunostained for surface receptors (an N-terminal antibody against GluA1 was used) before permeabilization for staining of intracellular antigens (MAP2). Coverslips were washed 3x with TBS and blocked for 1h at RT in TBS containing 5% BSA. Then, primary antibody (GluA1, see **Key Resources Table**) at 1:400 dilution was added to coverslips in TBS containing 1% BSA and incubated for 2h at RT or overnight at 4°C. Following 4X washes with TBS cells were incubated with 1:400 secondary, fluorophore-linked antibody (Alexa Fluor 488, cat. #: A-11034) in TBS containing 1% BSA. These steps were repeated to immunolabel MAP2 (1:400, see **Key Resources Table** for primary and secondary antibody information), with the addition of 0.1% Triton x- 100 in all solutions. Coverslips are then mounted on SuperFrost Plus slides in DAPI Vectamount medium.

## Acknowledgments

The authors thank Drs. Mark Dell’Acqua and Kevin Woolfrey (University of Colorado Denver) for the Akap79/150 antibody and NFAT-GFP constructs as well as constructive discussion, Dr. Sandra Hofmann (Univ. of Texas Southwestern) for the anti-PPT1 antibody, Dr. Froylan Calderon de Anda (Universitätsklinikum Hamburg-Eppendorf) for the GCaMP3 construct used to visualize calcium activity, and Dr. Daniel Dombeck (Northwestern Univ) for technical support and useful discussion regarding two-photon microscopy. This work is supported by startup funding awarded to A.Y. by the University of Illinois at Chicago, Department of Anatomy and Cell Biology and to S.M.C by the University of Illinois at Chicago, Department of Chemistry.

## Supplementary information

**Supplementary Video 1-related to Figure 4.** Live-cell calcium imaging of WT-vehicle treated cell treated with 1μm NASPM as indicated.

**Supplementary Video 2-related to Figure 4**. Live-cell calcium imaging of *Ppt1^-/-^*-vehicle treated cell treated with 1μm NASPM as indicated.

**Supplementary Video 3-related to Figure 4.** Live-cell calcium imaging of WT-TTX treated (1μm, 48h) cell treated with 1μm NASPM as indicated.

**Supplementary Video 4-related to Figure 4.** Live-cell calcium imaging of *Ppt1^-/-^*-TTX treated (1μm, 48h) cell treated with 1μm NASPM as indicated.

**Supplementary Video 5**. Representative spontaneous neuronal population activity in the visual cortex of an awake and behaving WT mouse using *in* vivo, two-photon calcium imaging.

**Supplementary Video 6.** Representative spontaneous neuronal population activity in the visual cortex of an awake and behaving *Ppt1^-/-^* mouse using *in* vivo, two-photon calcium imaging.

---

**Supplemental Figure S1-related to Figure 7.**
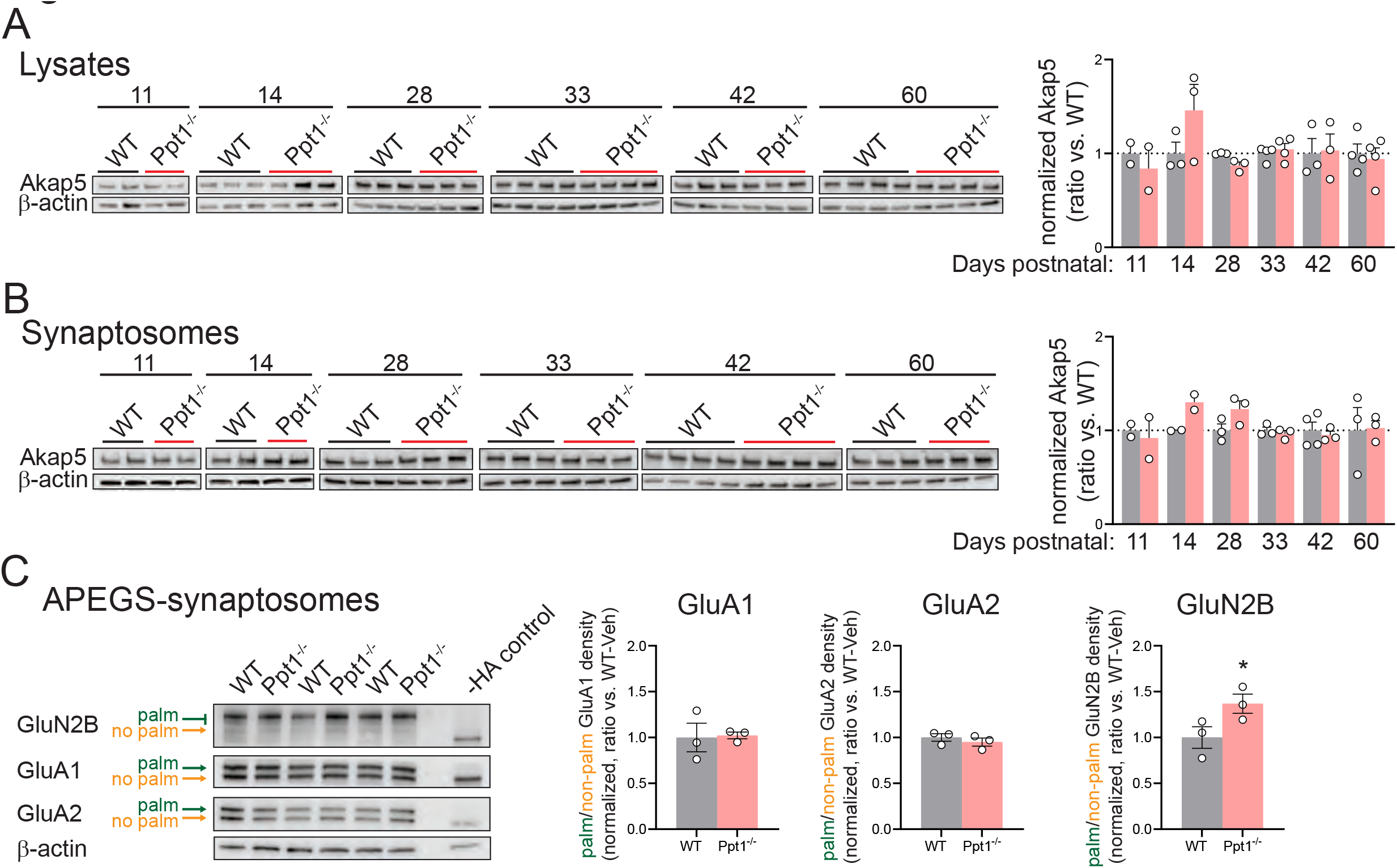
**(A)** Top ten enriched SynGO terms from proteins increased 1.2-fold in *Ppt1^-/-^* visual cortical lysates. **(B)** Network analysis of the genes increased in *Ppt1^-/-^* lysates by 1.2-fold that were annotated with the top biological process SynGO term “process at the synapse.”

**Supplemental Figure S2-related to Figure 7.**
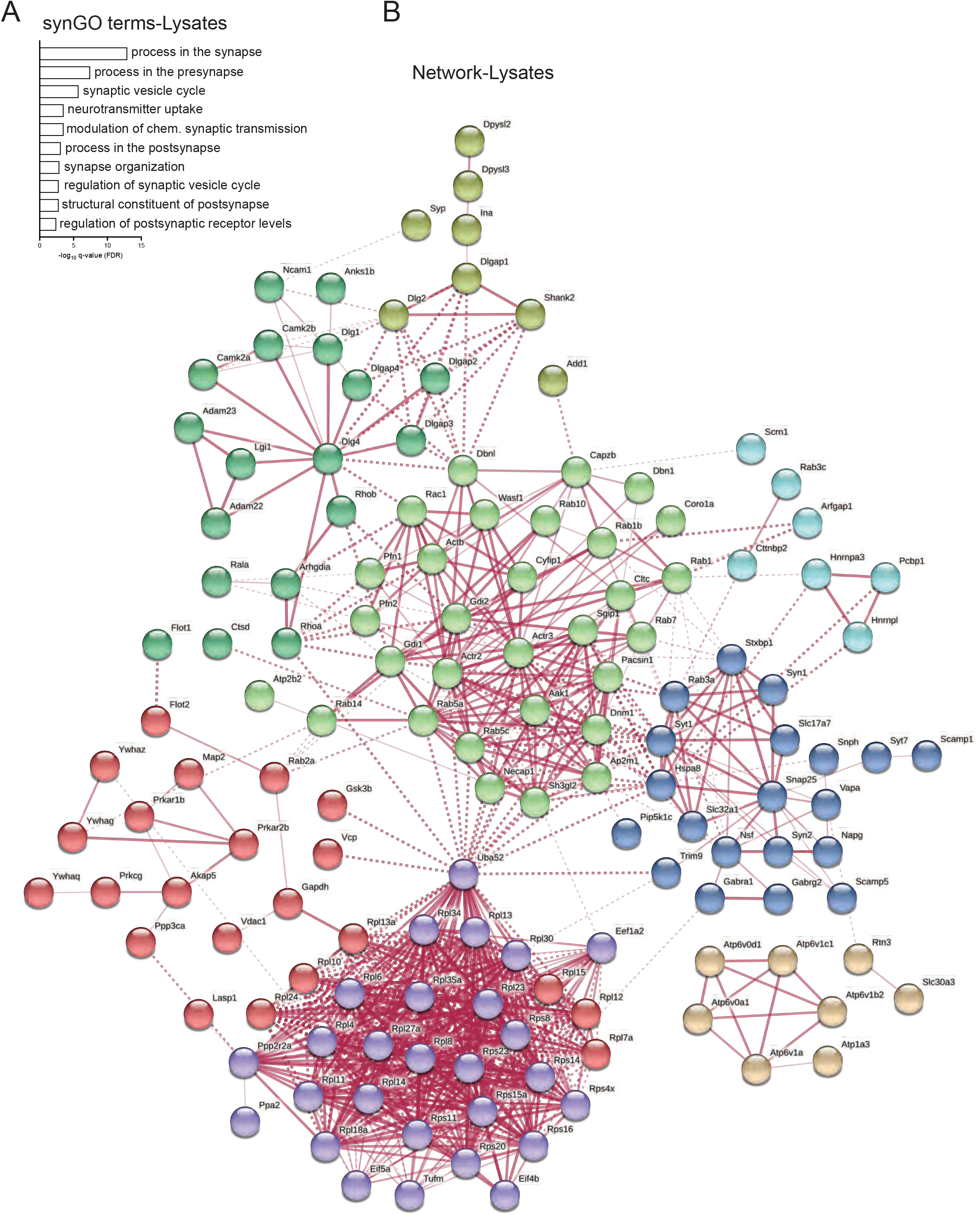
**(A)** Representative immunoblots (left) and quantification (right) of Akap5 levels in visual cortical lysates across age P11-P60. **(B)** Representative immunoblots (left) and quantification (right) of Akap5 levels in visual cortical synaptosomes across age P11-P60. **(C)** Representative blots of APEGS-processed synaptosomes (left) and quantification of the palmitoylated/non-palmitoylated ratio (right) of GluA1, GluA2, and GluN2B in visual cortical synaptosomes across age P11-P60. t-test: *p=0.0396. n=3 mice/group.

**Supplemental figure S3-related to Figure 8.**
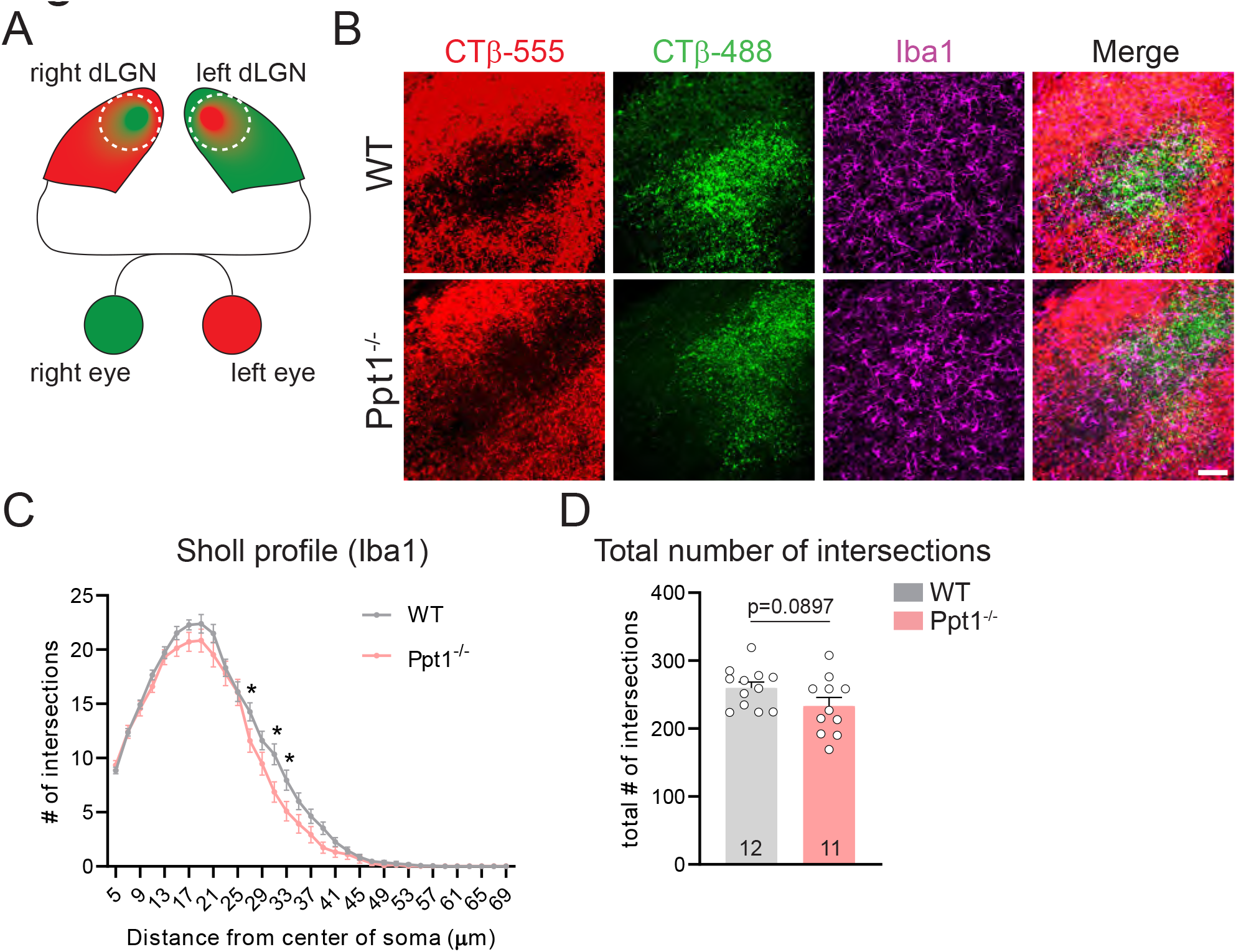
**(A)** Schematic of the injection paradigm for measurement of microglial morphology at the border between ipsilateral and contralateral retinogeniculate projections. **(B)** Representative images of CTβ 488, CTβ 555, and Iba1 immunostaining in the dLGN of a WT and *Ppt1^-/-^* mouse. Scale = 50μm. **(C)** Sholl analysis profile of microglia (Iba1) morphology in WT and *Ppt1^-/-^* dLGN between P21-42 (see **Table 2**). **(D)** Quantification of the total number of intersections/cell from Sholl analysis of microglia in the dLGN of WT and *Ppt1^-/-^* mice. N=10-11 animals/group.

**Supplemental figure S4.**
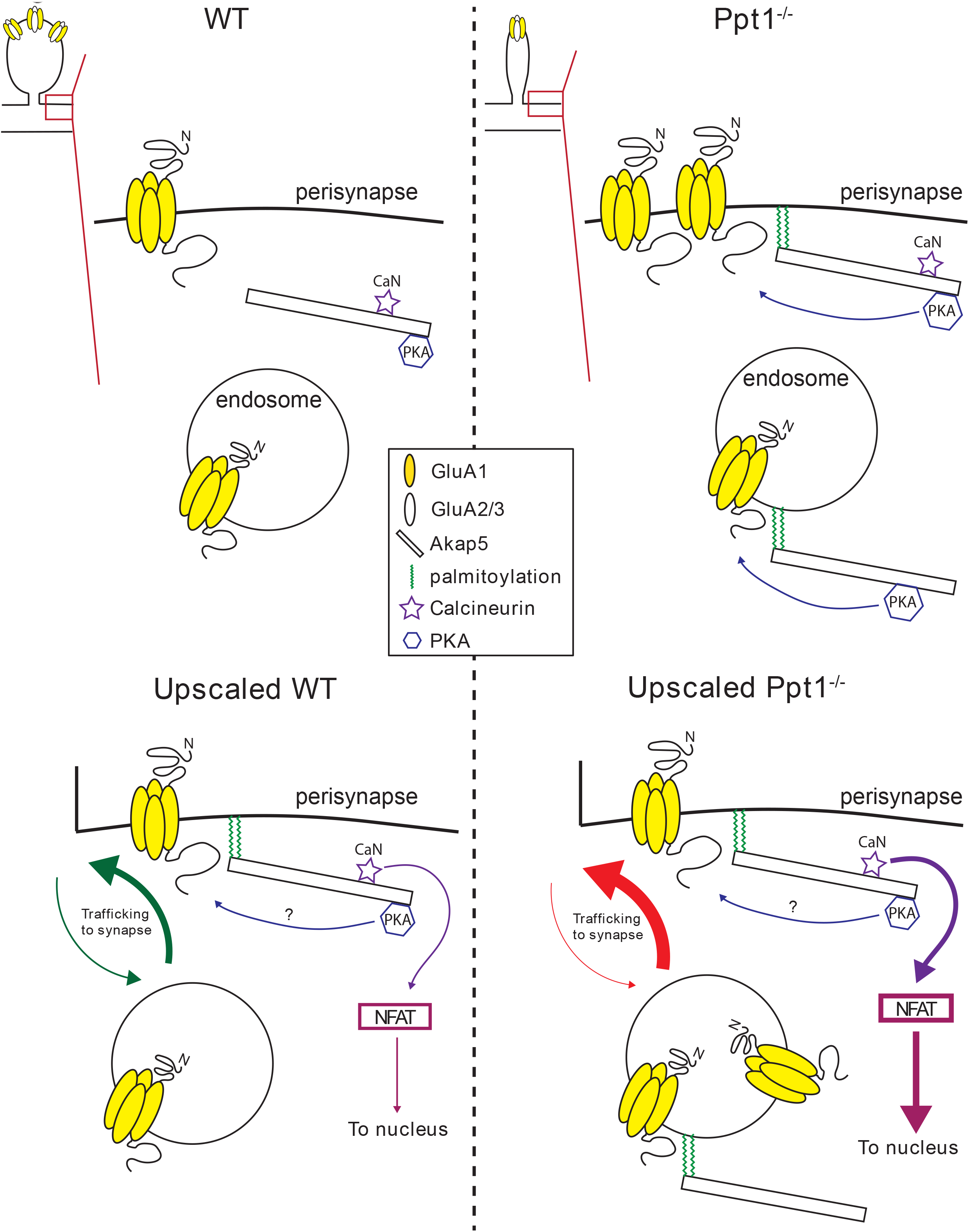
We propose that at baseline, where synapse morphology and composition are immature in *Ppt1^-/-^* neurons (top left), there is an abundant extrasynaptic pool of CP-AMPARs partly due to the over-palmitoylation of Akap5 (top). In response to the strong stimulus of synaptic scaling induction (bottom), perisynaptic CP- AMPARs are trafficked to the membrane of previously silent synapses, resulting in an exaggerated upscaling response that simultaneously drives enhanced proinflammatory signaling through the calcineurin-NFAT axis in *Ppt1^-/-^* neurons.

## Notes

### Competing Interest Statement

The authors have declared no competing interest.

